# Transplanted Autologous Neural Stem Cells Show Promise in Restoring Motor Function in Monkey Spinal Cord Injury

**DOI:** 10.1101/2023.05.06.539673

**Authors:** Razieh Jaberi, Reza Jabbari, Mostafa Hajinasrollah, Sara Mirsadeghi, Saeid Rahmani, Masoumeh Zarei-Khairabadi, Omidvar Rezaei, Seyed-Masoud Nabavi, Sahar Kiani

**Author notes:** Correspondence, (S.K.) Model.

## Abstract

Spinal cord injury (SCI) is a devastating condition that can result in permanent loss of motor and sensory function. In recent years, transplantation of neural stem cells (NSCs) has emerged as a promising therapeutic approach for SCI. However, adult NSCs residing in the central nervous system (CNS) show promise for cell-replacement therapy in neurodegenerative disorders. In this study, we aimed to investigate the efficacy of transplanting autologous NSCs isolated from the subventricular zone (SVZ-NSCs) into a spinal cord injury model in Rhesus monkeys. We induced SCI in eight Rhesus monkeys and then transplanted SVZ-NSCs into the injury site. Behavioral assessments and magnetic resonance imaging (MRI) were performed to evaluate the functional and structural recovery of the spinal cord. Histological analyses were also performed to assess the survival and differentiation of the transplanted cells. SVZ-NSCs were capable of expressing SOX2, Nestin, and GFAP in addition to self-renewing and spontaneous differentiation *in vitro*. The monkeys showed sensory and motor function recovery, spinal tract regeneration, and partial reconstruction of the spinal cord. Positive Reinforcement Training confirmed no adverse effects from the isolation procedure. Our findings suggest that NSCs transplantation could be a promising therapeutic approach for SCI in humans, and further studies are warranted to investigate its potential for clinical translation.

**Graphical abstract:** 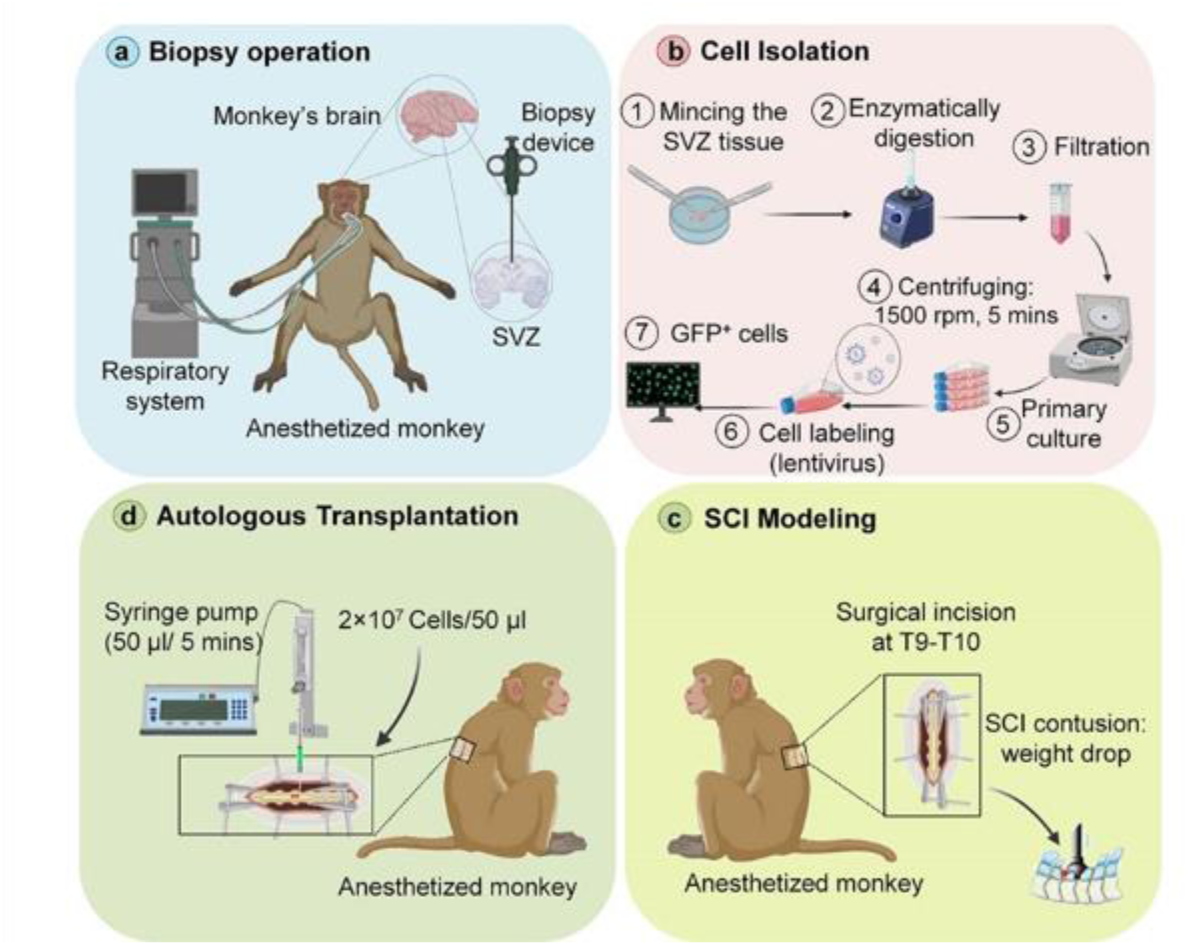

## 1. Introduction

Traumatic spinal cord injury (SCI) is considered the most devastating neurological condition to treat clinically [1], [2]. Based on statistics, three million people worldwide currently suffer from SCI, with approximately 180,000 new cases annually [3]. In fact, the prevalence of traumatic injury is 25.5/million/year in developing countries, having a far greater impact on the economic and social aspects of a patient’s life. Since 1700 BC, when SCI was described as an ’ailment not to be treated,’ no effective therapies have been identified [4] [5]. In humans, restoration of function after SCI remains an elusive goal, despite extensive advancements made in preclinical studies [6]. While common treatments such as medications, surgery, and rehabilitation provide effective amelioration in patients, none of these methods can compensate for cellular loss [7], [8]. Therefore, stem cell therapy has been identified as an appealing approach for cell replacement and axonal regeneration [9], [10].

Neural stem cells (NSCs) have the ability to promote cell replacement, axonal regeneration, and neural circuit reconstruction exclusively [11]. Furthermore, NSCs derived from various sources have already shown promising potential for SCI recovery in non-human and human primates [12]–[14]. Adult NSCs, as remaining reservoirs in the CNS, have long been discovered to substantially contribute to hippocampal neurogenesis during learning and memory, particularly in the human brain [15], [16]. In addition to their classical role in plasticity-associated neurogenesis, adult NSCs have been shown to be induced towards differentiation after brain impairments such as traumatic brain injury [17], ischemic brain injury [18], stroke [19], and epilepsy [20]. Adult NSCs have also been found to be capable of oligodendrogensis triggered by demyelination, in addition to neurogenesis [21]. Conversely, in several progressive disorders such as Alzheimer’s disease [22] and Parkinson’s disease [23], neurogenesis activation fails due to impaired circumstances. Similarly, stress and depression decrease neurogenesis while antidepressant agents can reverse this effect [24], [25].

The potential of adult NSCs to compensate for neuronal and glial loss has attracted significant attention in recent years, with numerous studies suggesting that targeting this source of NSCs holds promise for cell replacement therapies [26]–[28]. Promising approaches include managing adult NSCs’ migration paths in the brain [29], manipulating local cell types to create a permissive niche for neurogenesis [30], [31], and targeting adult NSCs via new-generation viruses to master their differentiation competency [32], [33]. However, a significant gap still exists between these methods and their application in the clinic. Recently, much research has focused on SVZ-NSCs, which hold multipotency, differentiation [34], and migration capacity [35]–[39]. Additionally, the differentiated cells derived from SVZ-NSCs are able to integrate into the pre-existing neural network. SVZ-NSCs are located within the walls of the lateral ventricles, which are accessible by penetrating non-eloquent parts of the human brain [40]. In fact, SVZ-NSCs are the ideal endogenous cell source for in vivo manipulation or harvesting for *in vitro* expansion and further autologous transplantation [41].

SVZ-NSCs are considered to be a promising candidate for cell-replacement therapy, not only for brain injuries but also for spinal cord injuries [42]. However, endogenous neural repair in the spinal cord is insufficient because the ependymal NSCs, which are the adult NSCs reservoir in the spinal cord, are unable to generate neurons. Moreover, the use of exogenous sources of NSCs is still controversial [43]. Therefore, in this study, we aimed to investigate the effectiveness of autologous SVZ-NSCs, which were grafted for the first time to a contusion SCI model in rhesus monkeys. As there are genetic, neurophysiological, and neuroanatomical similarities between humans and non-human primates [44], preclinical studies on lower primates are particularly important [13]. Additionally, unlike rodents, primates do not undergo significant auto-repair, similar to humans [45], [46], which is why monkeys are a privileged animal model in this study. Our hypothesis is that SVZ-NSCs can be used as a reliable cell source for SCI amelioration in primates, and this will overcome the sensory and motor dysfunction as well as cell rejection.

## 2. Materials and methods

### 2.1. Ethical Statement

The research procedures that involved primary cell culture and Neural Stem cells isolation and expansion, as well as animal studies (monkey brain endoscopy and spinal cord injury modeling), were conducted in accordance with the guidelines set forth by the “Royan Institute Ethics Committee” (IR.ACECR.ROYAN.REC.1397.104), (approval ID: EC/92/1009) (Figure S11) and also are presented according to ARRIVE guidelines [47].

### 2.2. Animals

Eight rhesus monkeys weighing between 3-6 Kg and aged between 3-6 years were obtained from the Primate Research Center of the ROYAN Institute. Prior to the study, the animals were carefully screened for intestinal parasites, tuberculosis, Simian Immunodeficiency Virus (SIV), herpes B, hepatitis A and B viruses, as previously mentioned (see Table 1). The monkeys were randomly divided into two groups (each animal was assigned a code prior to the division to blind all experimenters): the SVZ-NSCs Graft group (n=5) and the Control group, in which animals were injected with DPBS after SCI (n=3) (see Table 1). Depending on the site and severity of the traumatic injury, subsequent dysfunction can be categorized as partial or complete, specifically paraplegia or tetraplegia [48]. In this study, the SCI contusion model was induced at the T9-T10 spinal cord segment to induce further paraplegia (partial injury), and SVZ-NSCs were transplanted to the monkeys during the sub-acute phase. The animals were monitored for a total of six months from SCI through MRI; behavioral and electrophysiological evaluations were conducted, with the exception of histological analysis, which was only performed on subjects #06 and #07 after 9 and 10 MPI, respectively.

**Table 1.** Experimental Groups

| Subject | Injection | Amount of the cell transplanted | Weight before SCI (Kg) | Weight at the end of the study (Kg) | Control of the urinary drainage (W/MPI*) | Bedsore** | Muscle hindlimb atrophy | Survival time (month) |
| --- | --- | --- | --- | --- | --- | --- | --- | --- |
| #01 | PBS | 0 | 6.850 | 4.100 | Manually drainage up to end | Stage IV | Severe | 6 |
| #02 | NSC | $22 \times 10^6$ | 5.4 | 5.8 | 4 WPI | Stage III | Moderate | 6 |
| #03 | NSC | $21.2 \times 10^6$ | 6 | 7.35 | 2 WPI | Stage II | Mild | 6 |
| #04 | NSC | $23.5 \times 10^6$ | 6 | 7.1 | 3 WPI | Stage III | Mild | 6 |
| #05 | NSC | $21.4 \times 10^6$ | 5.8 | 6.31 | 2 WPI | Stage III | Mild | 6 |
| #06 | PBS | 0 | 6.68 | 3.67 | Manually drainage up to end | Stage IV | Severe | 9 |
| #07 | NSC | 24 × 10<sup>6</sup> | 6.3 | 6.750 | 3 WPI | Stage III | Mild | 10 |
| #08 | PBS | 0 | 6.47 | 4.9 | Manually drainage up to end | Stage IV | Severe | 6 |

\* W/MPI: Week or Month post injury
\*\* wound management: Ulcers were kept clean and the skin was dried, Zinc oxide ointment administration

### 2.3. SVZ-NSCs isolation

Animals were anesthetized with an intramuscular injection of ketamine (15 mg/kg) and xylazine (0.4 mg/kg). The monkey’s head was shaved and the skull was exposed, followed by a craniotomy to access the brain using a surgical drill. The monkey’s brain tissue was harvested from one SVZ (in each monkey) through neuro-endoscopy surgery with an endoscopic device (AUMAN medical, China) equipped with a high-resolution video camera to navigate and access the SVZ. The coordinate information was captured according to computed tomography (CT). A small piece (about 2 mm^2^) of the ventral wall of the SVZ was obtained from every monkey using biopsy forceps of the endoscopy instrument, in both the Graft and Control groups. After tissue isolation, connective tissue and skin were sutured, and the small superficial bones were replaced to the skull and connected by bone wax (MEGASPECTR, Bone wax 2.5G W810, Ukraine). The animals were then monitored and evaluated for normal locomotion and neurological index. The fresh tissues were transported in PBS (phosphate buffer saline) containing 10% penicillin-streptomycin (Sigma, P4333, USA) at 4°C, minced into small pieces, and digested enzymatically with Trypsin 0.25 mg/ml (Sigma, T4049, USA) and DNase I 80 U/ml (Sigma, D4527, USA). Centrifugation (1500 rpm for 3 minutes) was performed to remove the enzyme, and the cells were filtered using a nylon mesh with 70-micron pores (Corning, 431751, USA). The cells were then plated on T25 flask and incubated at 37°C/5% CO2 in neural stem cells medium (NSCM), which contained DMEM/F-12 supplemented with 0.05% B27, 1% N2, ITS (1 mg/ml insulin, 0.55 mg/ml transferrin, and 0.67 mg/ml selenium), 10% knockout serum replacement, 1% non-essential amino acids, 2 mM L-glutamine, penicillin 100U/ml and streptomycin 100 μg/ml (1%) (all from Invitrogen, USA), human basic fibroblast growth factor (bFGF, 40 ng/ml; ROYAN Institute), and Epidermal growth factor (EGF, 20 ng/ml; ROYAN Institute). The medium exchange was initiated after approximately nine days of the isolation concomitant with colony emergence in primary culture, followed by every other day medium replacement before passaging.

### 2.4. Immunocytochemistry staining and flow cytometry

After four *in vitro* passages, SVZ-NSCs were grown on 0.001% poly-L-ornithine (Sigma-Aldrich, P4707, the USA) and 10 mg/ml laminin (Sigma-Aldrich, L2020, the USA) seeded in 4-well dishes for further fixation and staining. SVZ-NSCs were fixed by paraformaldehyde (PFA) 4%, permeabilized by Triton 0.1% and then blocked by BSA (Sigma-Aldrich, A2058, the USA). The following primary antibodies were incubated with cells for an overnight at 4°C: anti-Nestin (1:200, Chemicon, MAB5326, the USA), anti-GFAP (1:400, Sigma-Aldrich, G3893, the USA), anti-SOX2 (1:200, Santa Cruz, Sc-20088, the USA), The next day, after the removal of primary antibody, the cells were incubated with either goat anti-mouse (1:1000, Invitrogen, A11005, the USA) or goat anti-rabbit (1:1000, Invitrogen, A11037, the USA) and incubated for 45 minutes at 37°C. The nuclei were stained via 4’, 6-diamidino-2-phenylindole (DAPI) (0.1 μg/ml, Sigma-Aldrich, D8417, the USA). Fluorescent imaging was performed using an Olympus IX71 Fluorescence Microscope (Olympus, Tokyo, Japan) with a DP72 digital camera. For analysis and cell counting, LS Starter 3.2 and ImageJ software were employed. For quantification, ten random fields were captured to count the cell nuclei (DAPI). Almost 1500 nuclei were calculated to find, and report expressed markers in percentage.

Flow cytometry was performed as the following: cells were collected enzymatically (trypsin) and then staining was performed with selective antibodies as following: anti-SOX2 (1:200, Santa Cruz, Sc-20088, the USA), and anti-GFAP (1:400, Sigma-Aldrich, G3893, the USA), Anti-Nestin (1:100, Chemicon, MAB5326, the USA), and anti-TUJ1 (1:400, Sigma-Aldrich, T8660, the USA), secondary fluorescent antibodies were anti-mouse IgG (1:200, Chemicon, AP308F, the USA) conjugated with FITC and FITC anti-rabbit IgG (1:200, Sigma-Aldrich, F1262, the USA). To detect and measure the positive cells, a BD FACSCalibur™ (India) instrument was employed, and histogram and other graphs were analyzed by <u>Flowing Software 2.5.1</u>.

### 2.5. SVZ-NSCs in vitro spontaneous differentiation

To address the multipotency of SVZ derived putative NSCs, the glial-derived neurotrophic factor (GDNF, 10 ng/ml (Sigma-Aldrich, G1777, the USA) and brain-derived neurotrophic factor (BDNF, 10ng/ml, Sigma-Aldrich, B3795, the USA) were substitute for both EGF and bFGF for two months to induce spontaneous differentiation in the cells. Thereafter, IF staining were performed for anti-TUJ1 (1:400, Sigma-Aldrich, T8660, the USA), anti-NG2 (1;200, Millipore, AB5320, the USA), anti-PDGFRα (1:200, Sigma-Aldrich, SAB4502142, the USA), Anti-Olig2 (1:100, abcam, ab109186, the USA), anti-GLAST (1:200, abcam, ab181036, the UK), anti-vGLUT1 (1:200, Abcam, ab134283, The USA), anti-NeuN (1:200, abcam, ab104224, the USA).

### 2.6. Colony formation assay

To investigate the self-renewal capacity of isolated SVZ-NSCs, a clonogenicity assay was performed. A diluted suspension of cells (100 cells/ml) was prepared and divided into a 96-well dish, with each chamber coated by 0.001% poly-L-ornithine (Sigma-Aldrich, P4707, the USA) and 10 mg/ml laminin (Sigma-Aldrich, L2020, the USA), containing 10 µl of the cell suspension. The cells were allowed to settle in the 96-well plate for one hour, after which 100 µl of neural expansion or neural stem cell medium (NSCM) was added to each well. Single cells were allowed to proliferate and enlarge in size for 21 days until their diameter exceeded 200 µm to investigate the colony formation capacity of SZV-NSCs. Colonies with the intended size were picked up to form neurospheres in the presence of neurosphere medium (NSM) containing 10% knockout serum replacement (KOSR).

### 2.7. Neurosphere Immunohistochemistry staining

Neurospheres were collected from NSM and fixed in PFA 4% overnight. Next, the PFA was removed, and the neurospheres were placed in 30% sucrose for about three hours. In the next step, the neurospheres were placed in a piece of agar gel, cut, and located in an aluminum foil frame exposed to droplets of the optimum cutting temperature glue (OCT Tissue-Tek® OCT™ Compound) slowly. The frozen blocks (−25°C to -30°C) were transferred to a cryostat (CM1950, Leica Biosystems, USA) for sectioning. Slices of 12 µm thickness were collected on glass coverslips and subjected to indirect immunofluorescence staining against CD133 (1:200, Abcam, ab19898, USA) as a surface marker, as described above.

### 2.8. Real-time qPCR analysis

Total RNA was manually extracted using TRIZOL reagent, and cDNA synthesis was performed using 1 µg of the total RNA (Fermentas kit, USA). To quantify real-time gene expression, the synthesized cDNAs were mixed with SYBR Green and QPCR Master Mix (Fermentas, USA), and the mixture was then placed inside a rotor gene 600 real-time PCR machine. The mRNA levels were normalized to GAPDH as an internal control. All the primers used were specific to humans and were previously implemented in our studies [34], [49].

### 2.9. Micro-electrode array recording

Early electrophysiological properties of differentiated SVZ-NSCs were assessed extracellularly using a micro-electrode array (MEA) system. All recordings were carried out inside a CO_2_ incubator (5%) on a hot stage (37°C) integrated with the amplifier (MEA1060-Inv, Multichannel systems, Germany). SVZ-NSCs autonomous differentiation was induced on a 60MEA200/10iR-ITO (Multichannel systems, Germany) under conditions similar to those mentioned in previous sections. Approximately 2×10^5^ cells were plated on a PLO substrate in differentiation medium. Signals were sampled at 25kHz and visualized via MC-Rack software (Multichannel systems, Germany). The spikes with frequencies higher than 300kHz were discriminated from total frequencies by bandpass (300 to 3000 kHz) Butterworth (order three) filtration, processed in R 4.0.3.

### 2.10. Contusive spinal cord injury model production

In this study, we induced an SCI contusion model in rhesus monkeys using a modified NYU impactor device, as previously described [49]. Briefly, the animals were anesthetized with intramuscular ketamine (15 mg/kg) and xylazine (0.4 mg/kg), and the spinal cord at T9-T10 was exposed by removing surrounding muscles, lamina, and spinous processes, while stabilizing clamps were used to secure the upper and lower spines (T8 and T11). A 50g rod weight was dropped vertically onto the spinal cord from a height of 12 cm, producing a contusion of approximately 10 mm^2^. Animals received twice-daily doses of 25 mg/kg cefazoline for one week following the injury. Daily bladder discharge was performed to assist with urinary retention, and if necessary, the animals were given Tramadol (20 mg/kg) for pain [50]. As per standard protocol for surgical procedures, the animals fasted for eight hours before and after the surgery (NPO), and Lactulose syrup was provided if needed to prevent stool retention.

### 2.11. Cell labeling and transplantation

To track the transplanted SVZ-NSCs in the injured monkey, we labeled isolated cells with GFP before injection. Lentivirus was utilized to deliver GFP into SVZ-NSCs prior to transplantation. Human embryonic kidney (HEK 293T) cells were transfected to produce lentivirus particles. Co-transfection of CMV-GFP construct and plasmids packaged with pCMV-vsvg and pCMV-gp was achieved using lipofectamine 3000 (Life Technologies). The medium of transfected cells was collected and concentrated by ultracentrifugation (20,000 for two hours at 4°C) to purify the released viruses. SVZ-NSCs were grown on tissue culture dishes coated with 0.001% poly-L-ornithine (Sigma-Aldrich, P4707) and 10 mg/mL laminin (Sigma-Aldrich, L2020). Cultured SVZ-NSCs were infected with GFP viruses only after the culture reached 70-80% confluency in NSC expansion medium (Figures 4B-G). As previously stated, two experimental groups were established: Graft and Control. After ten DPI, both groups of animals were anesthetized with intramuscular injections of ketamine (15 mg/kg) and xylazine (0.4 mg/kg). The spinal cord was then exposed again, under the same procedure as previously mentioned, for the cell injection. Graft animals were injected with 20 × 10^6^ GFP-labeled SVZ-NSCs suspended in DPBS (50 µL) using a Hamilton syringe (27G) and a micro-injector instrument (Stoelting, the USA) for 10 minutes [13] (Figure S4 C-D). Control animals underwent the same surgery procedure but received only DPBS without SVZ-NSCs.

### 2.12. Spinal cord histological evaluation

All animals were sacrificed for the histological analysis at the endpoint of this study, i.e., six MPI except for one Graft (survived until 10 MPI) and one Control (survived until 9 MPI). The fate of GFP-labeled SVZ-NSCs, horns and central canal structure were investigated by Immunofluorescence and H&E staining as mentioned in our previous study [49].

The spinal cord tissue was fixed in formalin sugar (30%) for preservation. The fixed tissues were then embedded in optimum cutting temperature (OCT) compound under freezing conditions of 25°C to -30°C using a Leica CM1850 instrument. Serial sectioning was performed transversely for four Graft and one Control animal, while sagittal sectioning was performed for one Graft and one Control animal. Ten series of 20 µm thick slices were collected on each slide to cover 200 µm of the spinal cord. The collected slices were then washed in PBS for 10 minutes, and permeabilized with Triton X-100 (0.5%) for 45 minutes to 1 hour depending on the target marker localization. Unspecific connections were blocked using a solution of 5% normal goat serum in PBS-T for 45 minutes at room temperature. After blocking, the sections were incubated with primary antibodies at 4°C overnight, including anti-GFAP, anti-NF200, anti-TUJ1, anti-DCX, anti-NG2, and anti-MBP (1:200, Santa cruz, sc-271524, USA). The primary antibodies were washed out with PBS-T, and the sections were then incubated with secondary antibodies (goat anti-mouse and goat anti-rabbit) at 37°C for 45 minutes. Finally, nuclei were stained with 4’,6-diamidino-2-phenylindole (DAPI) (0.1 μg/mL, Sigma-Aldrich, D8417). Fluorescent microscopy was performed using an OLYMPUS IX71 microscope. Cell counting in the immuno-stained sections was performed using ImageJ, where the images were converted into monochrome 8-bit, and the mean grey level number of black and white pixels was determined within the tissue.

To evaluate the anatomical structure of the spinal cord in both the Graft and Control groups, we performed serial sectioning on 2 cm of spinal cord and stained the sections using H&E (Figure S8 and S9). Briefly, the sections were hydrated three times in distilled H2O (3 min each), then immersed in a hematoxylin solution (GHS316, Sigma, Germany) for 4 minutes. After rinsing in warm running tap water (for 4 min), the slides were placed into lithium carbonate. Next, they were rinsed in distilled H2O (for 30 seconds) and in 95% reagent alcohol (for 30 seconds) before being counterstained with eosin Y solution (HT110116, Sigma, Germany) for 1 min. The sections were then dehydrated by placing them into 95% reagent alcohol twice, absolute reagent alcohol (100% EtOH) twice, and xylene (for 2 min each). Finally, we mounted the slides and captured images using a microscope (OLYMPUS, BX51, Japan).

### 2.13. Magnetic Resonance Imaging (MRI)

The visualization of neurological tissues, including the spinal cord, was performed using a 3-Tesla superconducting scanner (Siemens, Prisma, Germany) on sedated monkeys with ketamine (15 mg/Kg) and xylazine (0.4 mg/kg) at three different time points: Before injury (BI), to recognize the healthy anatomy of the animals; 48 hours Post Injury (PI), to confirm the contusion injury at T9-T10 level; and five Months Post Transplantation (MPT), to assess the impact of SVZ-NSCs on spinal construction. Sequences were obtained from sagittal and axial T1-weighted and T2-weighted MR scans between C5-L1 in all monkeys one week BI and 48 hours PI to confirm cavity formation as well as the absence of edema and hemorrhage [51].

### 2.14. Transcranial Magnetic Stimulation (TMS)

Transcranial Magnetic Stimulation (TMS) was used to assess hindlimb motor function using the MagPro X100 device (MagVenture, Denmark) at three time points: before injury (BI), post-injury (PI), and month days post-injury (MPI). To elicit a motor response, a circular coil (Cool-40, with transducer head dimensions of 52 x 54 x 42mm/ 2.0×2.1×1.7) was placed on the monkeys’ skull above the motor cortex. Animals were given ketamine (15 mg/Kg) and xylazine (0.4 mg/kg) to prevent unwanted movement during TMS. The superficial skin of the tibialis posterior muscle was shaved to locate and fix the record and reference electrodes, which were placed on the belly of the gastrocnemius muscle and the Achilles tendon, respectively. TMS was performed using a single stimulus with the following features: Time-based (5 ms/unit), sensitivity (200 µV/unit), lower frequency limit (20 Hz), and upper-frequency limit (2 kHz).

### 2.15. Walking corridor construction

To evaluate the walking ability, coordination, and basic learning capacity of monkeys, a walking corridor was constructed and utilized before biopsy, after biopsy, BI, PI, and after SVZ-NSCs transplantation. The transparent corridor, which was previously used in a similar study, was tailored to the body dimensions of the monkeys and consisted of three retractable segments, measuring 4.5 meters in length, 0.35 meters in width, and 0.75 meters in height. The animals were allowed to move freely through the corridor [51], and their reactions and movements were recorded by a digital camera (Nikon, D3500, Japan) during two tests that were implemented simultaneously in one experiment: the Monkey Hindlimb Score (MHS) and Positive Reinforcement Training (PRT). The camera enabled researchers to monitor the animals’ walking patterns, inner-step distances, standing tendencies and abilities, weight-bearing, and reward receiving for later analysis.

### 2.16. Behavioral, sensory, and motor evaluation

Sensory and motor functions were evaluated in both the Graft (n=5) and Control (n=3) groups up to six MPI. Sensory function was examined by evoking a response in the animals using a pinch stimulus to the toe, tail, and anus. The intensity of the response to the stimuli (graded from 0 to 3) was considered for toe and tail reflexes. The animal’s ability to respond to a physical pinch stimulus on the anal area was evaluated on a scale from 0 to 1.

To evaluate motor function in monkeys, we used the Monkey Hindlimb Score (MHS), which is a 9-point (0-8) scoring system (Table S2). To minimize bias, two blinded investigators watched videos of the monkeys’ movement in the walking corridor and assigned scores (0-9) for each hindlimb. We recorded videos of the monkeys moving freely in the corridor and calculated their improvement in motor ability as the mean percentage increase in hindlimb scores. To assess hindlimb engagement during standing and jumping, we conducted a climbing test by placing a banana above the monkeys’ living cage. Scores ranged from 0 for monkeys who did not attempt to climb to 100 for those who successfully climbed the rods and caught the banana. Monkeys who partially climbed the rods using only their hands or hindlimbs received a score between 0 and 100 (presented as a percentage). All data were presented as percentages.

### 2.17. Cognitive evaluation

To ensure that biopsy surgery did not have any unfavorable effects on animal’s basic cognitive abilities such as classical conditioning, we employed PRT before and after biopsy concomitant with MHS. For PRT, motivating foods were located at the end corner of the corridor to encourage the animals to take at least three independent steps. The test was established based on a reward given to the animals for walking along the corridor independently [52]. Three distinct conditions were considered for PRT in our study: 1. animals who did not take any step were not awarded any food, 2. animals who walked an incomplete path (to the middle of second segment) but did not reach the end of the corridor were awarded an apple, 3. successful animals that passed through all three segments to the end were awarded a banana. The number of times monkeys received rewards (apple or banana) was reported as a percentage. By performing PRT in the constructed corridor, we were able to encourage injured animals (PI) to take a step inside the corridor and walk through the path as long as they had the desire and strength to do so for motor function evaluations.

### 2.18. Statistics

The total numerical data, pertaining to all animals, are presented as mean ± SD of three independent experiments. The normality of the data was assessed by the Kolmogorov-Smirnov test. ANOVA and Tukey’s post hoc test were used for comparisons of means for SVZ-NSCs *in vitro* characterization (colony formation, positive cells in fluorescent imaging, gene expression), MTT staining, and IHC. Behavioral analyses (MHS, sensory evaluations, climbing test, and PRT) were assessed by the Mann-Whitney U test and unpaired t-test. A p-value of <0.05 was considered statistically significant. All data were analyzed using GraphPad Prism 8.0.1 software.

## 3. Results

### 3.1. Before Transplantation: In Vitro Characterization of Autologous SVZ-NSCs

Prior to transplantation of autologous SVZ-NSCs, after four *in vitro* passages (Figure 1A), we managed to determine the identity and capabilities of these cells from all aspects by immunofluorescent (IF) staining, flow cytometry, spontaneous differentiation, colony formation assay, qPCR, and microelectrode array (MEA) recording. SVZ-NSCs, from all monkeys, tended to form an extended spindle-like morphology with distinctive nuclei after five days of culturing. In the next ten days, they gradually became proliferative to generate cell aggregations (Figures 1B and B’ and S1 A). The proliferation rate, calculated by counting SVZ-NSCs in every four passages, showed a gradual increase between passages so that it reached the maximum rate at passage four (Figure 1C).

**Figure 1.**
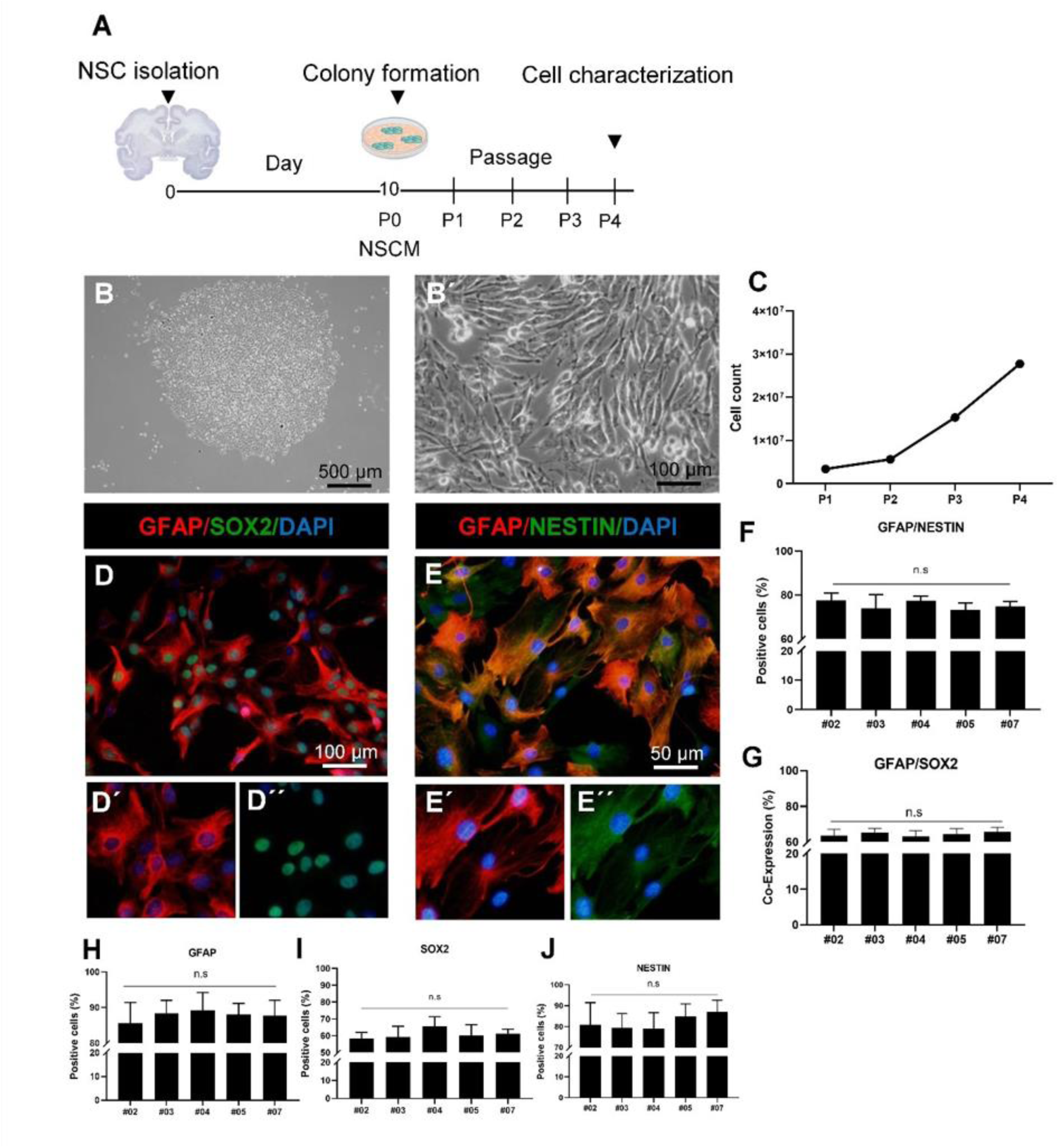
Primary culture and SVZ-NSCs *in vitro* characterization. **(A)** The cultural procedure for SVZ-NSCs is illustrated in a schematic diagram from isolation to passages. Briefly, SVZ cells were cultured until they formed distinct colonies in approximately five to ten days, and then passaged four times in NSC expansion medium (NSCM) before characterization. **(B)** Phase-contrast pictures show colony-forming SVZ-NSCs in primary culture and their morphological features at passage four. **(C)** SVZ-NSCs derived from monkey #03 proliferated normally, and their quantity enhancement was measured by carefully counting the cells at every passage, demonstrating their maximum quantity at passage four. **(D)** and **(E)** Immunofluorescence staining indicated that glial fibrillary acidic protein (GFAP) was co-expressed with transcription factor SOX2 (green) as well as with Nestin (green) in SVZ-NSCs isolated from monkey #03. In all figures, nuclei were stained with DAPI (blue). **(F)** and **(G)** The charts show the quantity of GFAP co-expression with Nestin and SOX2 in SVZ-NSCs of all five animals (Mean ± SD, p = 0.9971, one-way ANOVA, Tukey posttest). **(H)**, **(I)**, and **(J)** The quantified expression level of GFAP, SOX2, and Nestin in SVZ-NSCs (by cell calculation in ten random fields) was compared with no significant differences between the five Graft subjects (p = 0.9971, one-way ANOVA, Tukey posttest).

In addition to the proliferation rate, the protein expression of SVZ-NSCs was also carefully considered, especially for NSCs’ popular markers such as GFAP, SOX2, and Nestin. Glial fibrillary acidic protein (GFAP)-positive type B cells, resident in SVZ, have long been identified as adult NSCs [53] which, again here, SVZ-NSCs expressed GFAP around 80% (Figures 1D’ and 1E’ and S1 B). Transcription factor SOX2, existing in proliferating NSCs, was found to be expressed by SVZ-NSCs approximately 60% (Figures 1D’’ and S1 B). Nestin, as an intermediate filament protein, is always expressed by neural stem/progenitor cells in the intact brain [53]. According to our results, the SVZ-NSCs, isolated from all animals, expressed this marker by nearly 80% (Figures 1E’’ and 1F, S1 B).

Furthermore, to confirm the SVZ-NSCs’ capability of expressing dedicated proteins concurrently, co-expression of GFAP with SOX2 (Figure 1D-D’’ and 1G) as well as GFAP with Nestin (Figure 1E-E’’, 1F and S1) was determined by 85% and 90%, respectively. Cell characterization (protein expression) via IF technique was performed for isolated SVZ-NSCs from every individual subject of Graft and there were no significant differences between five subjects in the expression of these markers (p = 0.9971, one-way ANOVA, Tukey posttest) (Figure 1H-J).

Flow cytometry analysis was performed on SVZ-NSCs of each monkey to better confirm NSCs identity. SVZ-NSCs expressed GFAP in nearly 80%, Nestin above 70%, and SOX2 in roughly 50% of the cells (Figure S2 A). The expression of class III beta-tubulin (TUJ1) as a pre-mature, neuronal-committed cell indicator was also determined in less than 10% of the SVZ-NSCs (Figures S2 A). Moreover, the expression level of mentioned proteins in SVZ-NSCs derived from five Graft animals revealed no significant differences (p = 0.9992, one-way ANOVA, Tukey posttest) (Figure S2 B). In addition to mentioned above, we checked the chromosomal arrangement of the isolated cells for any undesirable chromosomal abnormality. The karyotype test illustrated normal arrangement and quantity of chromosomes in SVZ-NSCs (S2 C).

Eliminating the expansion growth factors bFGF and EGF from NSCM, except for BDNF and GDNF as basic components of the neural differentiation medium, suggests that isolated SVZ-NSCs can spontaneously differentiate into cells positive for TUJ1, NeuN (neuronal lineage), NG2, PDGFRα, Olig2 (oligodendrocyte lineage), and GLAST (astrocytic lineage) in a two-month period (Figures 2 A-H, S3 A). Our experiments revealed that the differentiated SVZ-NSCs are also able to express vGLUT1 at a rate of around 40% (Figures 2C’ and 2H, S3). SVZ-NSCs isolated from each Graft subject showed similar differentiation properties, and we did not observe significant differences in the expression level of TUJ1, GLAST, NeuN, vGlut1, and NG2 (p = 0.9992, one-way ANOVA, Tukey posttest) (Figure S3 B). An adult NSC must also self-renew to construct colonies or neurospheres *in vitro*. The self-renewal capacity of SVZ-NSCs was evaluated by assessing neurosphere generation, which, as expected, all five samples were capable of forming colonies (Figures 2I-K). We suspended the cells in a 96-well culture dish, and after 21 days, colonies with distinct borders appeared. Putative NSCs emanated homologous spheres with diameters of around 200 µm (Figure 2J). In the next step, SVZ-NSCs-derived neurospheres were characterized to be positive for the NSC-specific marker CD133 (Figure 2J). The quantification of colony organization of SVZ-NSCs was around 60% on average in all five source subjects, and there was no significant difference between these samples (p < 0.5, one-way ANOVA, Tukey posttest) (Figure 2K). To further examine the stemness trait of SVZ-NSCs, RT-PCR was performed for several markers, namely GFAP, Nestin, SOX2, and PAX6. Among these markers, GFAP and Nestin showed higher expression than SOX2 and PAX6, respectively, and no significant differences were detected between subjects compared to the internal control (GAPDH) (p < 0.0001, one-way ANOVA, Tukey posttest) (Figure 2L). Moreover, MEA recording during SVZ-NSCs autonomous differentiation indicated that spike-generating neurons existed in the cell population even in the first month of induction (Figure 3).

**Figure 2.**
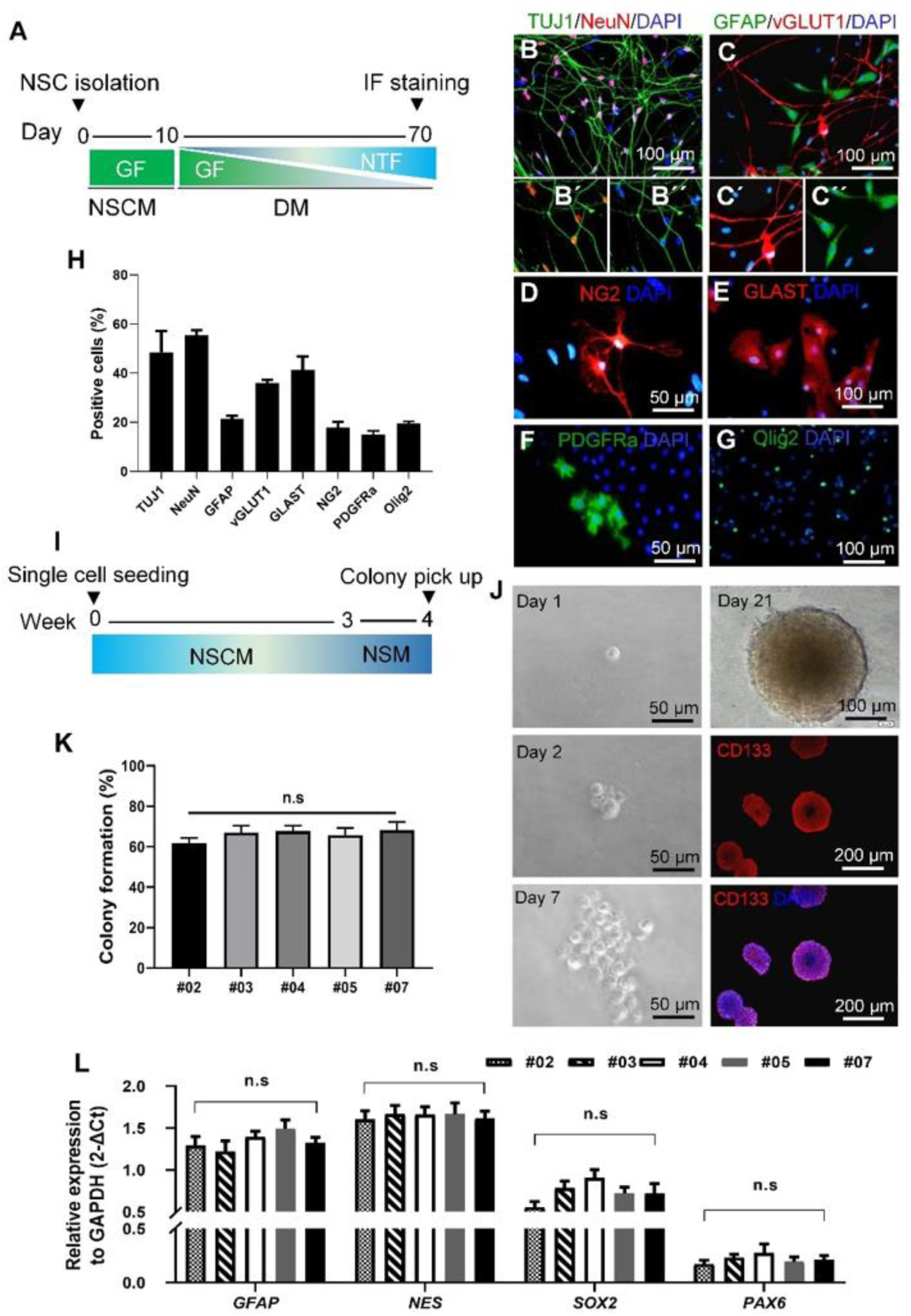
Self-renewal, multipotency and gene expression in SVZ-NSCs. **(A)** The diagram presents a schematic representation of the process of spontaneous differentiation of SVZ-NSCs (GF: Growth Factor, NTF: Neurotrophic Factor, NSCM: Neural Stem Cell Medium, and DM: Differentiation Medium). **(B-G)** During spontaneous differentiation of SVZ-NSCs in the #03 subject, TUJ1 (green), NeuN (red), and vGLUT1 (red) were expressed as neuronal markers, while GFAP (green), GLAST (red), NG2 (red), PDGFRa (green), and Olig2 (green) were expressed as glial markers, demonstrating the tripotency of these cells. Nuclei were stained with DAPI (blue). **(H)** The quantification graph, generated from 10 randomly chosen fields, illustrates the percentage of positive neuronal and glial markers after autonomous differentiation of SVZ-NSCs derived from the #03 monkey. **(I)** The schematic diagram outlines the colony formation assay procedure, in which single SVZ-NSCs were seeded in NSCM to form colonies in 21 days, and then, the organized colonies were picked up one week later (NSM: Neurosphere Medium). **(J)** Representative phase-contrast images reveal the time sequence of colony formation in the #03 subject, starting from 1, 2, 14, and 21 days after SVZ-NSC plating. Additionally, the colonies expressed CD133, an NSC marker. **(K)** The quantification of the colony formation capability of SVZ-NSCs, isolated from five diverse monkeys, did not indicate any significant differences (p = 0.3584, one-way ANOVA, Tukey posttest). **(L)** Quantitative real-time PCR was performed to assess the expression levels of SOX2, a stemness gene, as well as Nestin, GFAP, and PAX6, which are additional NSC markers in SVZ-NSCs at passage four. The analysis revealed no significant differences between cells isolated from distinct monkeys (p < 0.0001, two-way ANOVA, Tukey posttest).

**Figure 3.**
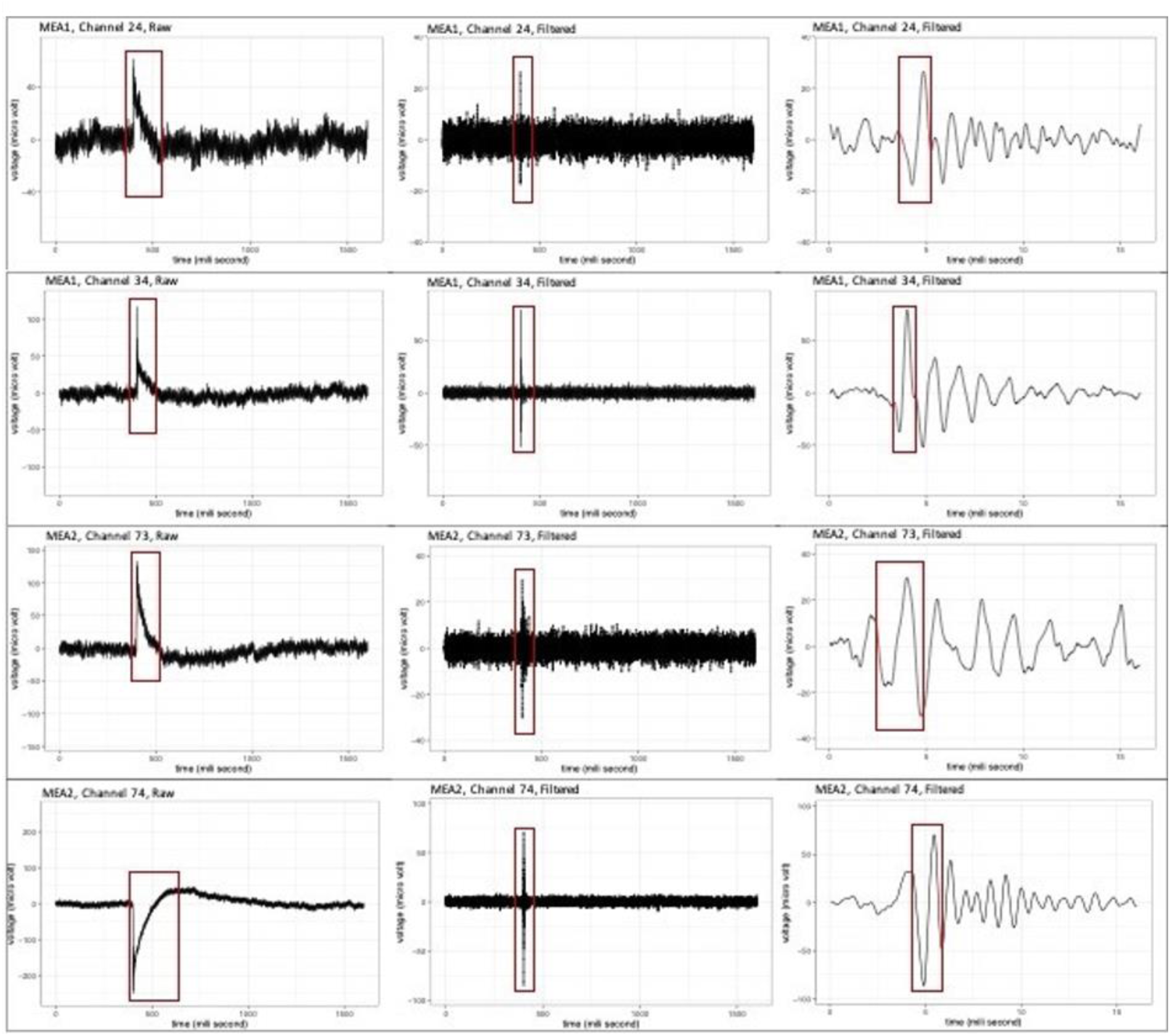
Early electrophysiological properties of differentiating SVZ-NSCs. Random active electrodes, comprising the spontaneous electrical activity of differentiating SVZ-NSCs on MEA, were depicted in three different temporal scales. **(A)** In the raw data, a complex signal containing low and high-frequency components of cell activity was detected. **(B)** A Butterworth-order 3 filtration discriminates spikes, corresponding to membrane action potential, owning frequencies higher than 300 kHz from other low-frequency components of the signal. Also, Fourier transform better uncovers the real amplitude of the filtered data, ranging approximately between -85 to 80 µv. **(C)** As illustrated in magnified filtered data, the action potential’s positive and negative waves are translated in reverse when recorded extracellularly.

### 3.2. SVZ-NSCs reconstructed the spinal cord structure

Sagittal MRI measurements were taken at different time points, including before injury (BI), 48 hours post-injury (PI), and five months PI, for all monkeys. Representative T2-weighted sagittal images of the undamaged spinal cord, as well as DTI at BI in the thoracic level, showed the normal anatomy of the spinal cord and regular cerebrospinal fluid (CSF) density (Figure 4A and S4 A-B). The midline incision’s surgical effect was evident in the MRI results at the contusion site 48 hours PI, with the MRI showing that the spinal cord was properly crushed at the T9-T10 level at the same time points (Figure 4A). Additionally, the injured site was visually recognizable (S4 C) until the SVZ-NSCs transplantation (S4 D). Five months PI, the partially reconstructed spinal cord was visible in the altered T2 intensity of Graft subjects, indicating the positive impact of SVZ-NSCs on the anatomy amelioration of SCI compared to Control (Figure 4A).

**Figure 4.**
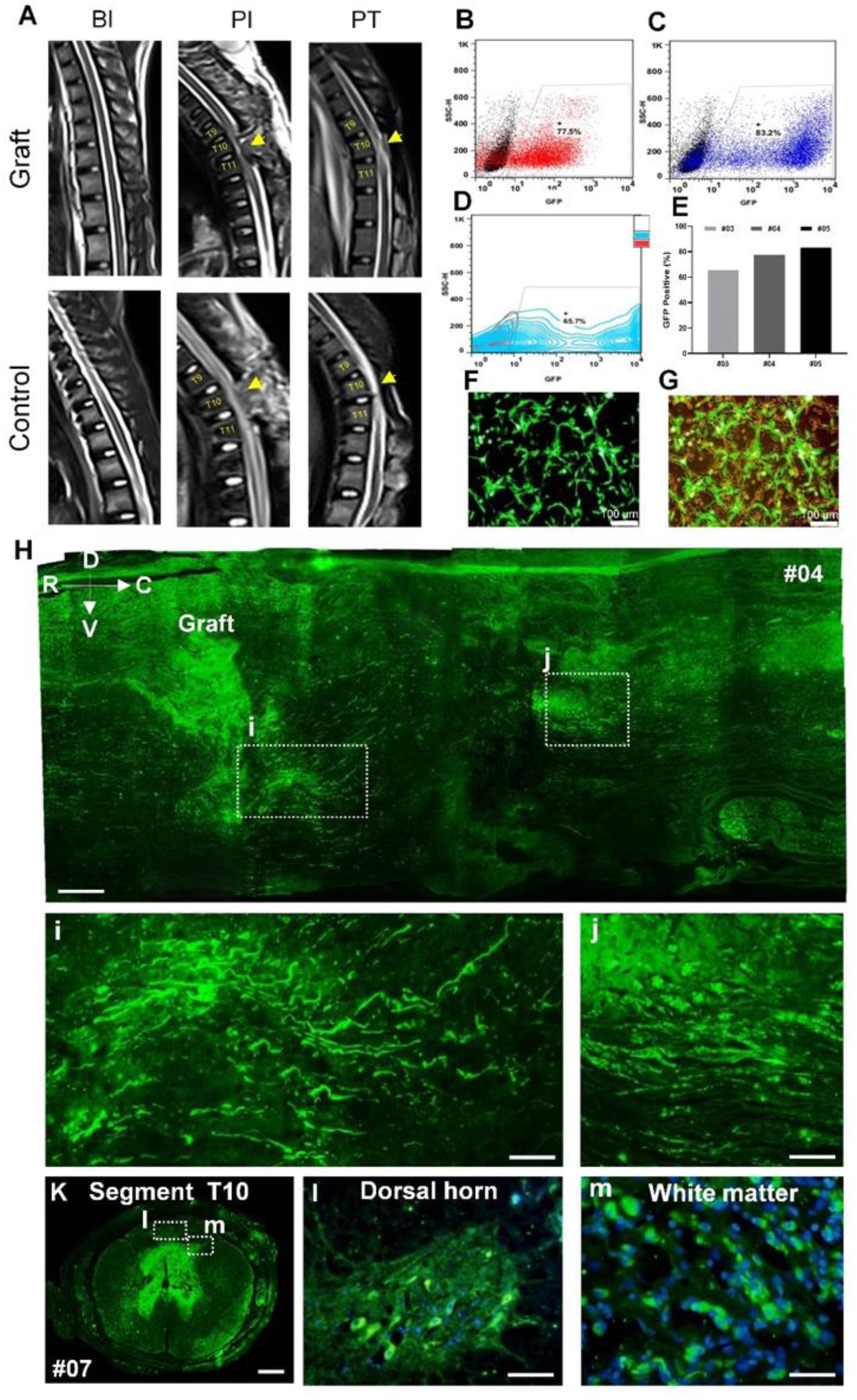
MR imaging, cell labeling and tracking. **(A)** The T2-weighted MRI signal depicts the normal anatomy of the spinal cord before injury and the disrupted structure of the spinal cord after SCI in the #03 monkey. Additionally, the MRI reveals partial reconstitution after six months in the Graft group compared to the Control group. **(B-G)** The efficiency of SVZ-NSCs transfection with GFP was measured by flow cytometry and visually observed under a microscope, with more than 60% of cells showing positivity for GFP. **(H-J)** A distinct population of transplanted SVZ-NSCs-GFP was detected rostral and caudal to the injury site six months post-injury in the #04 monkey. **(K-M)** GFP+ cells were also observed at the dorsal horn and white matter of the T9-T10 segment in the #07 monkey ten months post-injury.

### 3.3. Surviving SVZ-NSCs generate neurons and glia at the site of injury

To monitor the transplanted SVZ-NSCs, they were marked with GFP, as mentioned in the methodology. The effectiveness of transfection was evaluated by flow cytometry in three different cell populations prior to transplantation, demonstrating that roughly 70% of the cells were successfully transfected by the GFP vector. The GFP+ cells were observed to have survived and were found in both the rostral and caudal directions of the lesion site as well as in the gray and white matter of the T9-T10 segment of sagittal and transverse sections of the spinal cord, at six and ten MPI, as illustrated in Figures 4B-G and 4H-M, and S4 E-G.

Sensory and motor functions are severely affected by the loss of neurons and myelinating oligodendrocytes after injury. The formation of synapses and reconstruction of neuronal networks is critical for the accurate operation of these functions. Cell-based therapy is used to achieve this goal. In this study, we evaluated the in vivo fate of transplanted SVZ-NSCs (GFP+ cells) by assessing the expression of neuronal and glial markers as well as neuroblast and progenitor markers to determine whether they participated in synapse formation. DCX labeling revealed that less than 15% of the SVZ-NSCs remained undifferentiated after six MPI (Figure 5B 1 and C), while GFAP-GFP+ cells were detectable by around 20% (Figure 5B 2 and 5C). TUJ1 and NF200 expression suggested that SVZ-NSCs could differentiate into mature neurons *in vitro* (Figure 5B 3-4 and 5C), while vGLUT1 expression was found in 40% of cells after six MPI (Figure 5B 5 and 5C). The oligodendrocyte lineage was observed by almost 20% expression of NG2 and MBP (Figure 5B 6, 7 and 5C). These results indicate that SVZ-NSCs primarily differentiate into neuronal and oligodendrocyte lineage cells, which may contribute to functional improvement.

**Figure 5.**
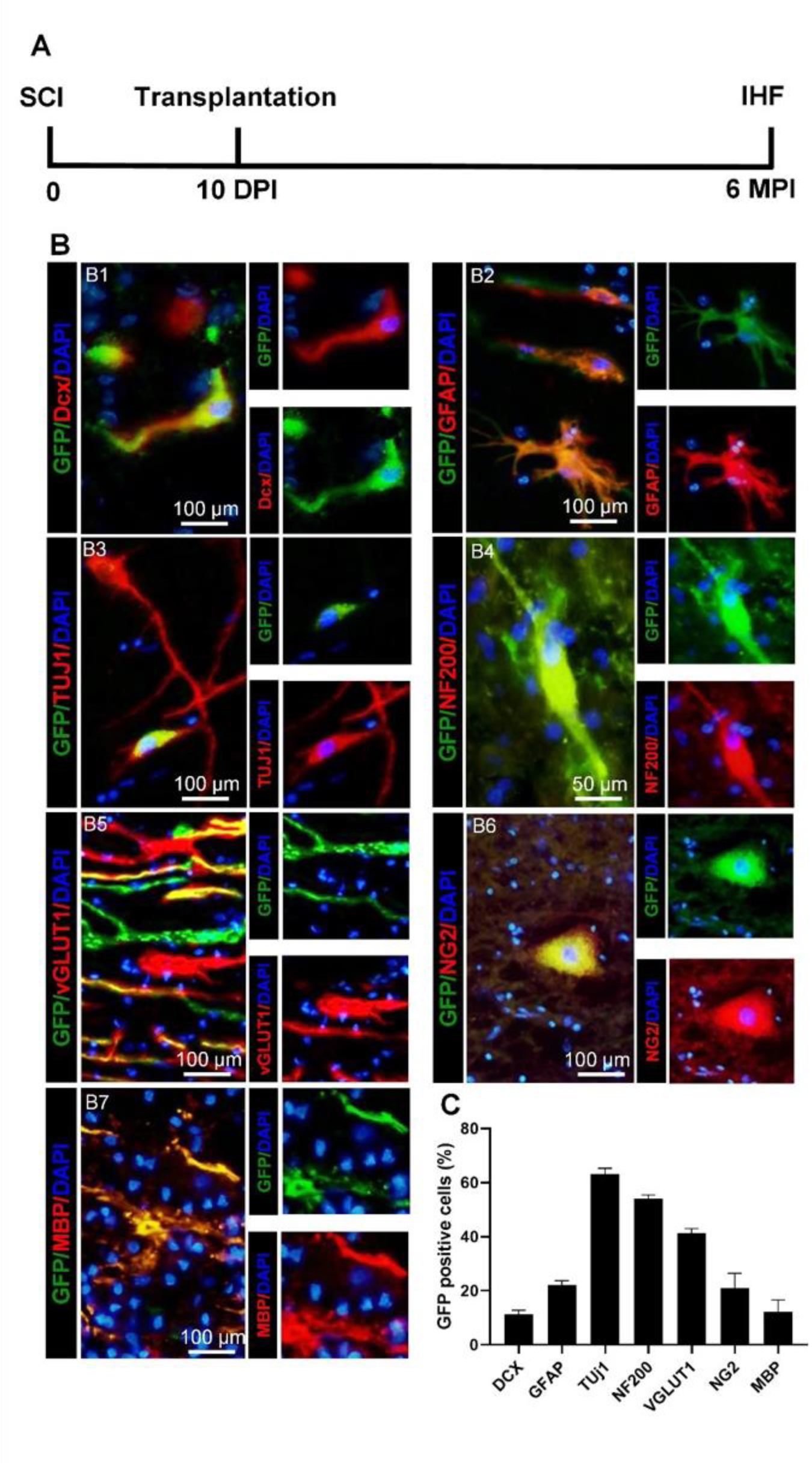
SVZ-NSCs differentiation into neuronal and glial cells after autologous transplantation. **(A)** The transplantation of SVZ-NSCs was performed 10 days after injury in accordance with the road map. Immunofluorescence staining was performed after six MPI for all subjects, except for #07, who was sacrificed at 10 MPI. **(B)** The expression of GFP was observed in conjunction with various markers: **(B1)** DCX for neuroblasts, **(B2)** GFAP for glial progenitor cells, **(B3-B4)** TUJ1 and NF200 for neurons, **(B5)** vGLUT1 for synaptic representation, **(B6)** NG2 for oligodendrocyte progenitors, and **(B7)** MBP for myelination. These markers were evaluated in #03 monkey six MPI after transplantation. **(C)** The quantified results of SVZ-NSCs progenies in #03 monkey’s spinal cord showed that the majority of GFP+ cells that survived differentiated into neurons (TUJ1+, NF200+, and vGLUT1+) and oligodendrocyte progenies (NG2+ and MBP+), while some cells remained undifferentiated (DCX) or in a progenitor state (GFAP).

### 3.4. Autologous transplantation of SVZ-NSCs improves sensory and motor recovery in SCI monkeys

In both the Graft and Control cases, sensory and motor function were blindly scored weekly during the first two months after the surgery, and evaluations were performed monthly thereafter until six months. After one-month post-injury (MPI), the sensory function returned to baseline in the Graft group. After three MPI, sensory recovery was observed in this group and remained at a plateau until the end of the study (six MPI). In contrast, animals in the Control group showed stable sensory disability without any significant recovery until the fourth MPI. Overall, toe and tail reflex demonstrated a considerable enhancement; however, anal reflex elevation was not significant (p < 0.05, Mann-Whitney U test) (Figures 6A-C, S5 A-C΄). Grafted animals responded to tail pinch faster than those in the Control group and had more powerful movement in response to tail pinch. After five MPI, animals in the Graft group could ultimately move their tails voluntarily.

**Figure 6.**
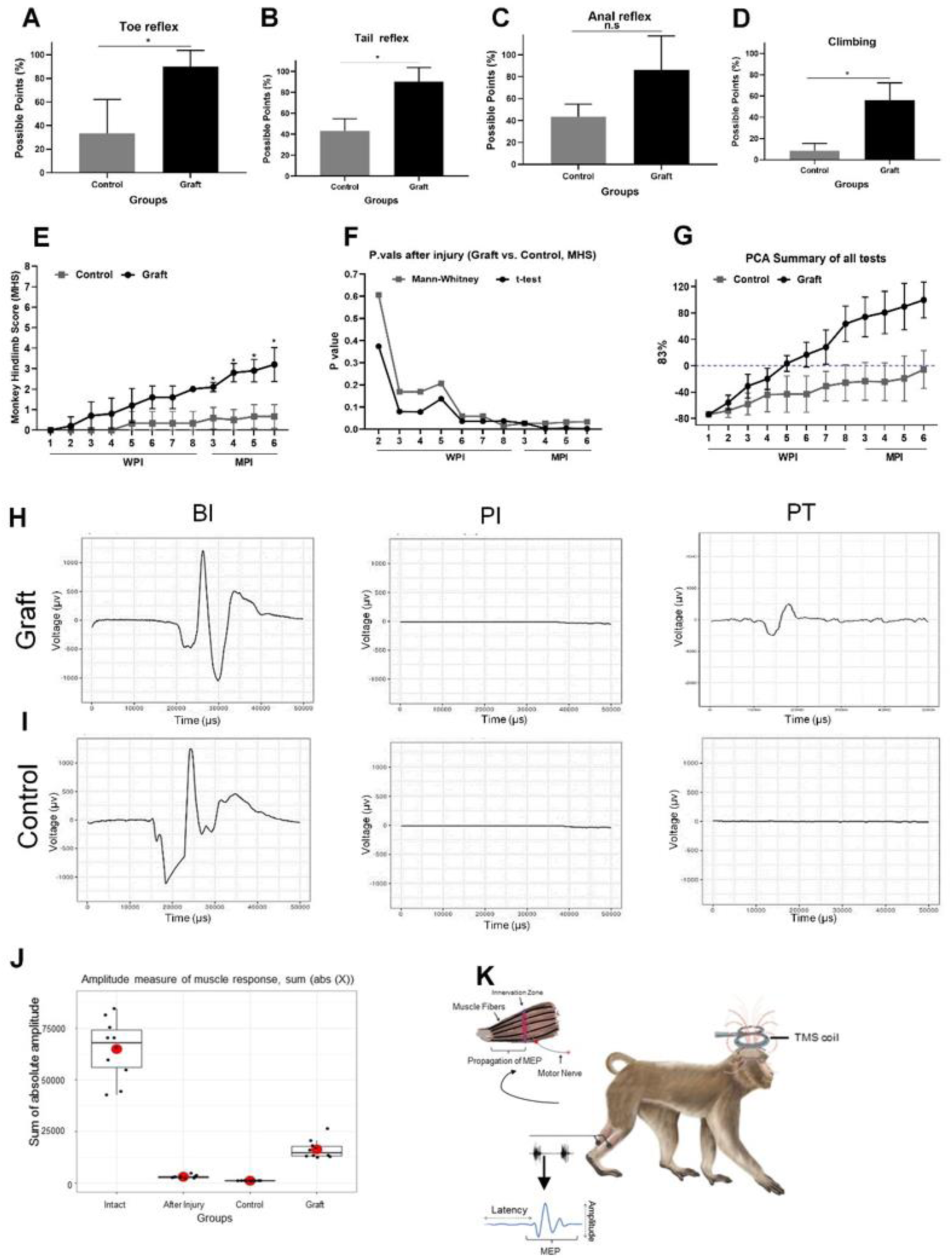
Behavioral evaluations and TMS outcome. **(A)** and **(B)** Sensory responses; toe and tail execution were significantly improved in the Graft in comparison to Control group (*p* < 0.05, Mann-Whitney U test), however, **(C)** anal reflex ability did not change significantly in between Graft and Control (*p* < 0.05, Mann-Whitney U test). **(D)** The ability of the animals to climb the cage bars were also elevated in Graft as shown in the column graph (*p* < 0.05, Mann-Whitney U test). **(E)** Overall performance of the monkeys, evaluated by MHS, was increased dramatically during six MPI and showed remarkable enhancement especially in the last four months in Graft comparing to the Control group (n=3) (*p* < 0.05, Mann-Whitney U test). **(F)** Non-parametric (Mann-Whitney U test) and parametric evaluation is represented for MHS during the follow up. (**G)** PC1 visualizes all behavioral assessment results for each monkey over six months in which Graft shows intense separation from the Control (WPI; Week Post Injury, MPI; Month Post Injury). (**H)** and **(I)** A common baseline evoked signal was observed in muscle of the Graft and Control BI. 48 hours following SCI, the muscle responses were eliminated in all monkeys, in consequent to cortical TMS. However, the muscular contraction signals were relatively returned in Graft in six MPI while we could not observe any response in the control animals. (**J)** Absolute amplitude, recorded at diverse timepoints of pre-and post-injury in Graft and Control groups, showed considerable differences between these two groups after six MPI (*p* < 0.0001,). **(K)** Graphical view of the TMS (TMS: Transcranial Magnetic Stimulation) and muscle recording in monkeys, schematically illustrated that the TMS coil were positioned on motor cortex.

To assess hindlimb locomotion in monkeys, we used the 8-point monkey hindlimb score (MHS), which evaluates movement in the hip, knee, and ankle joints. These three joints are essential for hindlimb locomotion in monkeys and are therefore important indicators of overall locomotor function.

Prior to injury, both groups of intact monkeys scored the maximum of eight on the MHS, indicating their ability to walk and run independently and change direction to reach rewards in the testing corridor, as trained by PRT. However, after a single week post-injury (WPI), all animals in both groups scored zero due to their inability to move their hindlimbs and joints, forcing them to rely on their forelimbs to pull their body forward to reach rewards in the corridor. By two weeks post-injury (MPI), monkeys in the Graft group were able to wiggle three hindlimb joints for sitting and standing, resulting in more favorable MHS scores than the Control group. The grafted monkeys were able to move considerably more easily than the Control animals, which only exhibited perceptible movement in one or two hindlimb joints. Additionally, the Graft group showed a significant improvement in their climbing ability after six MPI (p < 0.05, Mann-Whitney U test), as demonstrated by the overall percentage of effort exerted by the animals to climb the rods and reach the banana reward (Figure 6D, S5 D-D΄).

Our data show that transplantation of SVZ-NSCs resulted in significant locomotor recovery in the injured animals after six months post-injury. These animals were able to move more joints efficiently and showed some effort in standing (p < 0.05, Mann-Whitney U test) (Figure 6E, F and S6).

To visualize the locomotor function and sensory responses in each individual monkey over time, we used linear principal component analysis (PCA) and pooled the data. Our analysis revealed that PC1, representing the pooled data for each time point in the Graft group, significantly differed (elevated) from the Control group over time (Figure 6G). These results suggest that auto-transplantation of adult NSCs resulted in much more desirable functional and behavioral improvements in SCI monkeys at six months post-injury.

### 3.5. SVZ-NSCs are sufficient to improve evoked muscular responses through spinal tracts

Muscular contractions evoked by motor cortex TMS were recorded from animals in both groups at three time points: before injury (BI), post injury (PI), and six months post injury (six MPI). Baseline normal muscle responses were observed in all rhesus monkeys in both groups. Following contusive SCI, the muscle response was markedly abolished in both Graft and Control groups, but it returned to baseline in the presence of SVZ-NSCs at six MPI. In contrast, in the Control monkeys, there was no remarkable improvement in muscle response after SCI, even at six MPI with DPBS injection (Figure 6H-I). We measured the sum of absolute amplitude at each time point in both groups, and pooled the results across different animals at the BI and PI time points since the data were similar. After cell transplantation, the amplitude significantly differed from that in control animals (p < 0.0001, unpaired t-test) (Figure 6J-K).

### 3.6. The endoscopic biopsy itself did not impose any cognitive side effect to monkeys

To determine the safety of endoscopy surgery and its potential for damage to the surrounding tissues of the SVZ, we performed PRTs. Both groups of subjects quickly learned the task execution rules and were able to receive rewards through positive reinforcement while avoiding non-rewarded performances. Our data indicated that the endoscopy surgery did not adversely affect the learning ability of any of the subjects. In fact, they displayed an even greater level of expertise after the endoscopy, similar to what is expected from healthy animals (P < 0.05, paired t-test) (Figure S7 A). Furthermore, MRI results showed normal SVZ anatomy and its surroundings after biopsy (S7 D and E).

## 4. Discussion

In this study, we propose a translational application for autologous adult NSCs obtained from SVZ in a primate (rhesus monkey) model of SCI. The accessibility of SVZ for obtaining autologous NSCs, combined with the similarity of monkey and human spinal cord anatomy and pathophysiology, makes them a preferred model for this study. SCI results in permanent deficits in monkeys that resemble those seen in humans [45], [46], making them an ideal surrogate for evaluating the efficacy of autologous NSCs in improving SCI. In accordance with ethical considerations, we limited the number of animals and addressed key questions about biopsy surgery and the effects of SVZ-NSCs on monkeys by dividing them into only two experimental groups (Graft and Control).

In our previous report [34], we identified the *in vitro* properties of monkey SVZ-NSCs. In this study, we aim to compare various SVZ-NSCs derived from different individuals before engraftment. Our *in vitro* characterization results showed that SVZ-NSCs co-expressed GFAP with Nestin by almost 90% and GFAP with SOX2 by around 60%, regardless of the monkey source, consistent with previous reports [53]. The relative gene expression levels of GFAP, Nestin, SOX2, and PAX6 were also similar between samples, indicating that SVZ-NSCs have equal properties regardless of the animal source. The self-renewal capacity of SVZ-NSCs was confirmed to be analogous between all five sources using colony formation assays. SVZ-NSCs, like all NSCs, have the ability to spontaneously differentiate into three neural cell lineages (neuronal and glial). The spontaneous differentiation capability of SVZ-NSCs was examined in a BDNF/GDNF-containing medium, and TUJ1/NeuN co-expressing neurons, in addition to vGLUT1+/GFAP+ cells, accompanied by GLAST, NG2, Olig2, and PDGFRa-positive cells in the differentiated population, reflected the tripotency identity of all SVZ-NSC samples. The medium was formulated to avoid nudging the cells in a specific direction for maturation and only to address their intrinsic potency. Further *in vitro* maturation of oligodendrocyte progenitor cells into MBP+ or PLC+ oligodendrocytes requires the administration of particular growth factors such as IGF-1, NT3, and T3 [54]. Extracellular recording during the first month of *in vitro* spontaneous differentiation showed the emergence of single spikes in several electrodes, suggesting heterogeneous fate decisions between individual SVZ-NSCs, starting early after bFGF and EGF elimination. The appearance of higher frequency spikes (higher than 300kHz), corresponding to membrane action potential [55], demonstrated that a population of SVZ-NSCs can differentiate into electrophysiologically active cells in a permissive (not directed) circumstance. Further dedicated manipulation of this cell source *in vitro* would be beneficial to uncover their potency in producing distinct functional neurons, but in this study, we focused on identifying their preliminary features as reliable NSCs. Based on our investigations, SVZ-NSCs isolated from all five monkeys were evaluated with similar properties.

My apologies for the confusion earlier. Here’s the revised paragraph with grammar and clarity checks:

We conducted a thorough review of literature to determine the most effective vector and promoter combination for achieving long-term, sustainable, and non-targeted GFP expression in monkey-derived transplanted SVZ-NSCs, following our comparative *in vitro* identification of these cells. Lenti-viruses have been shown to provide long-lasting gene expression in both proliferating and post-mitotic cells, without any cytopathic effects [56], [57]. They are also known to be suitable for *in vivo* investigations of the CNS [58] and have been used in a two-year clinical trial for gene delivery to patients with Parkinson’s disease [59]. Furthermore, they have been employed for labeling hNSCs for traceability in rat spinal cord [60]. The CMV promoter has also been suggested as a reliable means for *in vivo* tracing of adult NSCs [61], [62], and it has been found to be optimal for *in vivo* transduction of both neuronal and glial cells in monkeys [63]. Therefore, we used a lentivirus construct to package the CMV-GFP to ensure detection of the triple progeny of SVZ-NSCs further MPI.

The study began by selecting monkeys with normal spinal cord anatomy, as determined by MRI and DTI scans. Contusion SCI was induced in these monkeys using a standard surgical procedure, which involved dropping a weight of 50 grams from a distance of 12 cm, as previously described in our publication [49]. MRI imaging confirmed the successful induction of SCI at the T9-T10 level, where the spinal cord was damaged and the dura was disrupted. The loss of muscle response to TMS at 48 hours post-injury further confirmed the SCI modeling. Subsequently, we observed that the GFP+ SVZ-NSCs survived and matured over time, with the majority of cells dedicated to neuronal fate, as quantified by *in vivo* tracing, equal with embryonic NSCs [13]. We detected GFP+/TUJ1+, GFP+/NF200+, and GFP+/vGLUT1+ cells six months post-injury. Furthermore, we found that the second largest population of differentiated SVZ-NSCs was dedicated to oligodendrocyte progeny, as indicated by the presence of GFP+/NG2+ and myelinating GFP+/MBP+ cells. A minority of SVZ-NSCs remained undifferentiated, at the neuroblast (GFP+/DCX+) and progenitor (GFP+/GFAP+) stages.

The differentiation of SVZ-NSCs into neurons and myelinating oligodendrocytes may potentially compensate for the cellular loss in spinal cord circuits and underlie the onset of functional recovery in subjects. To assess the improvement in spinal tract function following SVZ-NSCs transplantation, TMS was utilized [64], [65]. When compared to the baseline recording of intact animals, spinal cord injured subjects did not exhibit any response to cortical TMS. However, during the six MPI, a distinct response of approximately ± 200 µV was observed in recipient animals, evoked by cortical TMS. Furthermore, the prolonged absence of a reaction in Control and IF results indicates that SVZ-NSCs maturation could play a crucial role in Graft animals’ recovery.

Following SVZ-NSCs transplantation, an aroused movement (limb withdrawal, licking or touching) was observed in Graft subsequent to tail and limb pinch reflecting the sensory response recovery in animals. Furthermore, a withdrawal reflex has been found in the hind limbs in response to a deep pain stimulation (needle stimulus). Animals also responded to stimulus by receding their body from the needle rather rapidly. There was an improvement in both groups in clinical outcomes, but in the Graft, amelioration occurred much earlier than in Control. The induced injury on T9-T10 level caused immobility in hindlimbs joints (hip, knee, or ankle) and this deficiency was permanent in Control animals over time (measured by MHS). As opposed to Control, SVZ-NSCs induced perceptible movement of three joints in Graft subjects, though, without weight-bearing. Elicited from other preclinical reports and our own results, it seems that the mitigation in corticospinal tracts (according to TMS data) has led to the improved behavioral outcome (motor function and climbing ability) in Graft group here as well [66]. Some studies focused on the rehabilitation associated with cell therapy for enhancing functional recovery [67]. In fact, we also believe that controlling the muscle atrophy by rehabilitation in contribution with neuronal replacement in the spinal tracts, will bring premiere consequences for patients in movement and function. In our case, SVZ-NSCs seem to be a reliable cell source to repair the spinal tracts for exploiting an improved motor behavior in SC injured animals. In addition to behavioral analysis, MRI depicted that the spinal cord anatomy was recovered partially following the SVZ-NSCs autograft. This visualized data proved that spinal cord anatomy repair is associated with SVZ-NSCs *in vivo* differentiation in addition to behavioral and muscular response improvement.

NSCs were isolated through endoscopy, which is a minimally invasive technique [40], [68]–[71], from the SVZ. During the procedure, the surgeon carefully passed through the non-eloquent structures of the brain to reach the SVZ tissue, aiming to prevent any unfavorable damage. To do so, the endoscopic device was directed slowly to the brain from the longitudinal fissure, the most recognizable hollow structure of the brain.

After passing through the corpus callosum, the device reached the ventral wall of the SVZ, allowing for a small biopsy to be harvested. Despite the surgeon’s proficiency and caution during the operation, we thoroughly examined the animals for any possible side effects on cognitive abilities or coordinated movement resulting from the biopsy, considering that the limbic system, responsible for conditional learning, is located above the SVZ, and the caudate nucleus, in charge of coordinated movement, is located right beneath the SVZ [72]. Based on the MRI results, the endoscopic surgery was performed properly. In addition to the MRI, PRT was chosen to test if the animals could still reinforce a learned task after the biopsy [73], [74]. Monkeys were trained to walk along a corridor by placing a banana at the end corner, allowing for the MHS to be performed concurrently. Our findings suggested that the animals were able to remember and execute the task considerably better over time after the SVZ-NSCs autograft. To ensure the safety of the animals, their behavior was monitored by a camera inside the corridor and by a researcher frequently inside their living cage to detect any signs of uncoordinated movement after the biopsy. The monkeys were able to stand, walk and jump coordinately in a normal manner after the biopsy surgery and BI. Overall, our RPT and eye-monitoring results indicate that the endoscopy was safe throughout the duration of the study.

The application of NSCs for the treatment of SCI in humans has been proposed for a long time [75], and it is currently undergoing evaluation through three clinical trial phases, continuing until 2022 [76], highlighting the value of this cell type. Although different phases of SCI are considered for therapy in clinical trials, in the present preclinical examination, we preferred the sub-acute phase, which occurs between 10 to almost 30 days after SCI, to deliver the SVZ-NSCs dressing as soon as possible to the injury site, bypassing at least the primary severe inflammatory adverse effects on SVZ-NSCs survival [77]. According to our results, an approximately 15-day *in vitro* expansion period was determined to be sufficient to attain a relevant quantity of SVZ-NSCs for autograft during the sub-acute phase of SCI. Comprehensive good manufacturing practice (GMP) protocols for NSCs *in vitro* expansion also suggest substantial yield and viability of the cells during an equivalent period [78]–[80]. Hence, the selection of the sub-acute phase is entirely reasonable to achieve both an adequate SVZ-NSCs quantity for transplantation and impactful in situ operation, even for future clinical trials.

## Author contributions

Razieh J. performed cell culture, colony formation assays, Immunostainings, flow cytometry, RT-PCRs, spontaneous differentiation, cell preparation for autograft, behavioral assessments, histological analysis, figure preparation, data collection, statistical analysis, drafted the manuscript and participated in experimental design. Reza J. harvested SVZ tissue via biopsy in addition to SCI modeling. Mostafa H., as the veterinarian surgeon, performed SCI modeling, *in vivo* cell transplantation and animal caring and clinical assessments. Sara M. performed MEA recording, contributed to writing the paper and revised the scientific content of the reported results (as a blind reviewer). Saeid R. performed the statistics; MEA and TMS analysis. Masoumeh Z. K. labeled the cells by viral vectors, wrote the corresponding section in manuscript and contributed to histological analysis. Omidvar R. provided the clinical facility for surgery and animal care. Seyed-Masoud N. contributed to manuscript writing and submitting. Sahar K. was the main innovator of the proposed hypothesis, supervised the whole project, discovered the grant sources and contributed to manuscript writing and editing.

## Funding

This project was financially supported by grants from the ROYAN Institute, and the Iranian Council of Lotus (ICL).

### Institutional Review Board Statement

The study was conducted according to the guidelines of the “ROYAN Institute Ethics Committee” (IR.ACECR.ROYAN.REC.1397.104).

### Data Availability Statement

All data supporting the findings of this study are available within the article and its supplemental data file or from the corresponding author upon reasonable request.

## Supporting information

Figure S1: Characterization of Isolated Cells from Monkeys: Analysis of Morphological Properties and Neural Stem Cell Markers, Figure S2: fellow cytom

## Acknowledgments

We would like to express our sincere gratitude to thank Prof. Hossein Baharvand, Head of the Stem Cell Biology and Technology Department, for his support and also ROYAN animal facility center.

## Abbreviations

SCI: Spinal Cord Injury
NSCs: Neural Sem Cells
CNS: Central Nervous System
SVZ-NSCs: NSCs isolated from the Subventricular Zone
MRI: Magnetic Resonance Imaging
CT: computed tomography
PBS: Phosphate Buffer Saline
NSCM: Neural Stem Cells Medium
NSM: Neurosphere Medium
bFGF: basic Fibroblast Growth Factor
EGF: Epidermal Growth Factor
PFA: Paraformaldehyde
KOSR: Knockout Serum Replacement
OCT: Optimum Cutting Temperature glue
MEA: Micro-Electrode Array
MPI: Month Post Injury
DAPI: 4’,6-Diamidino-2-Phenylindole
BI: Before Injury
PI: Post Injury
PT: Post Transplantation
WPI: Week Post Injury
MPT: Months Post Transplantation
TMS: Transcranial Magnetic Stimulation
MHS: Monkey Hindlimb Score
PRT: Positive Reinforcement Training.

