## Supplementary material for "Transplanted Autologous Neural Stem Cells Show Promise in Restoring Motor Function in Monkey Spinal Cord Injury": Figure S1: Characterization of Isolated Cells from Monkeys: Analysis of Morphological Properties and Neural Stem Cell Markers, Figure S2: fellow cytom: Supplementary files.pdf

Fig. S-1

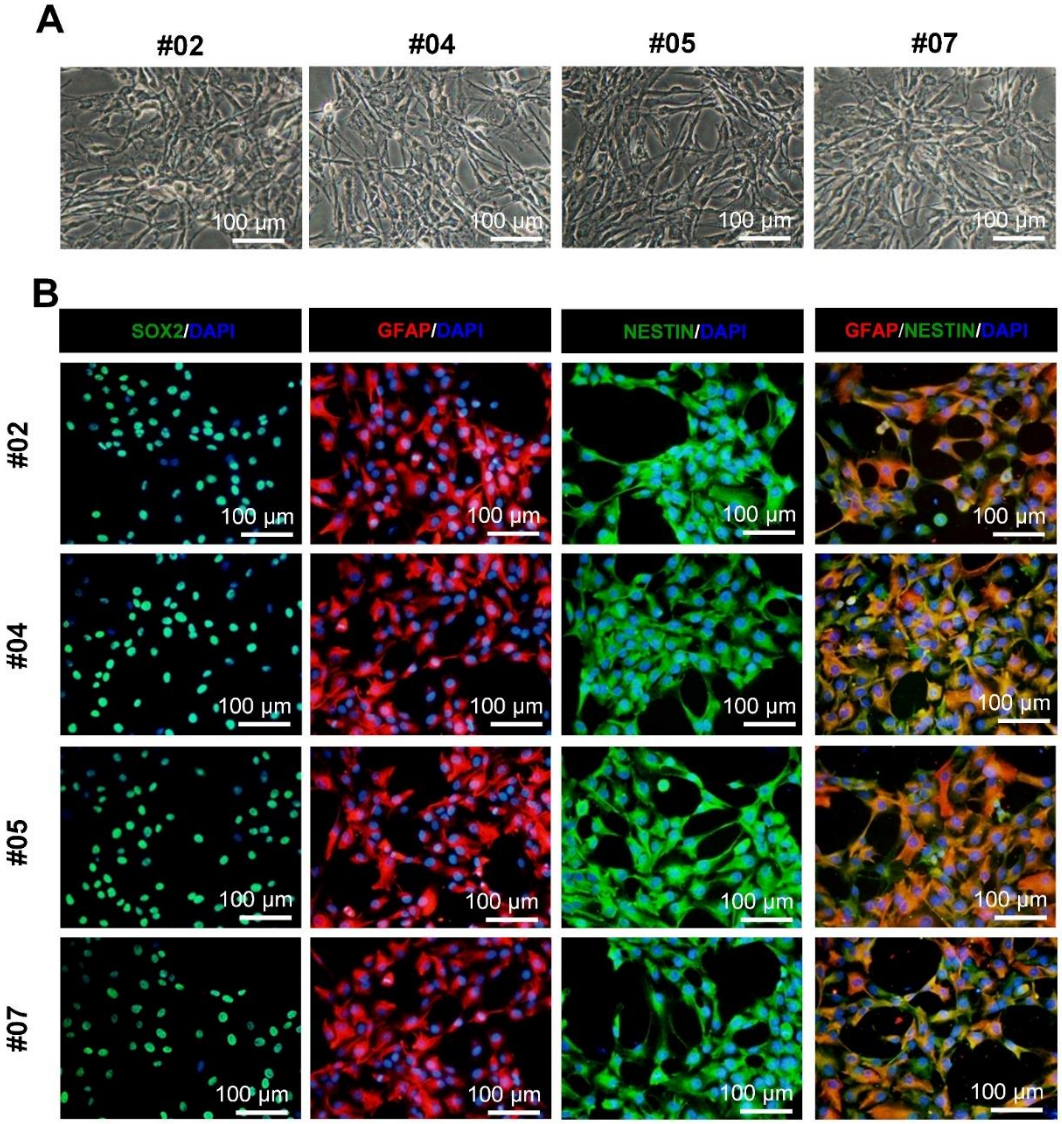


**Figure S1: Characterization of Isolated Cells from Monkeys: Analysis of Morphological Properties and Neural Stem Cell Markers**

**Fig. S-2**

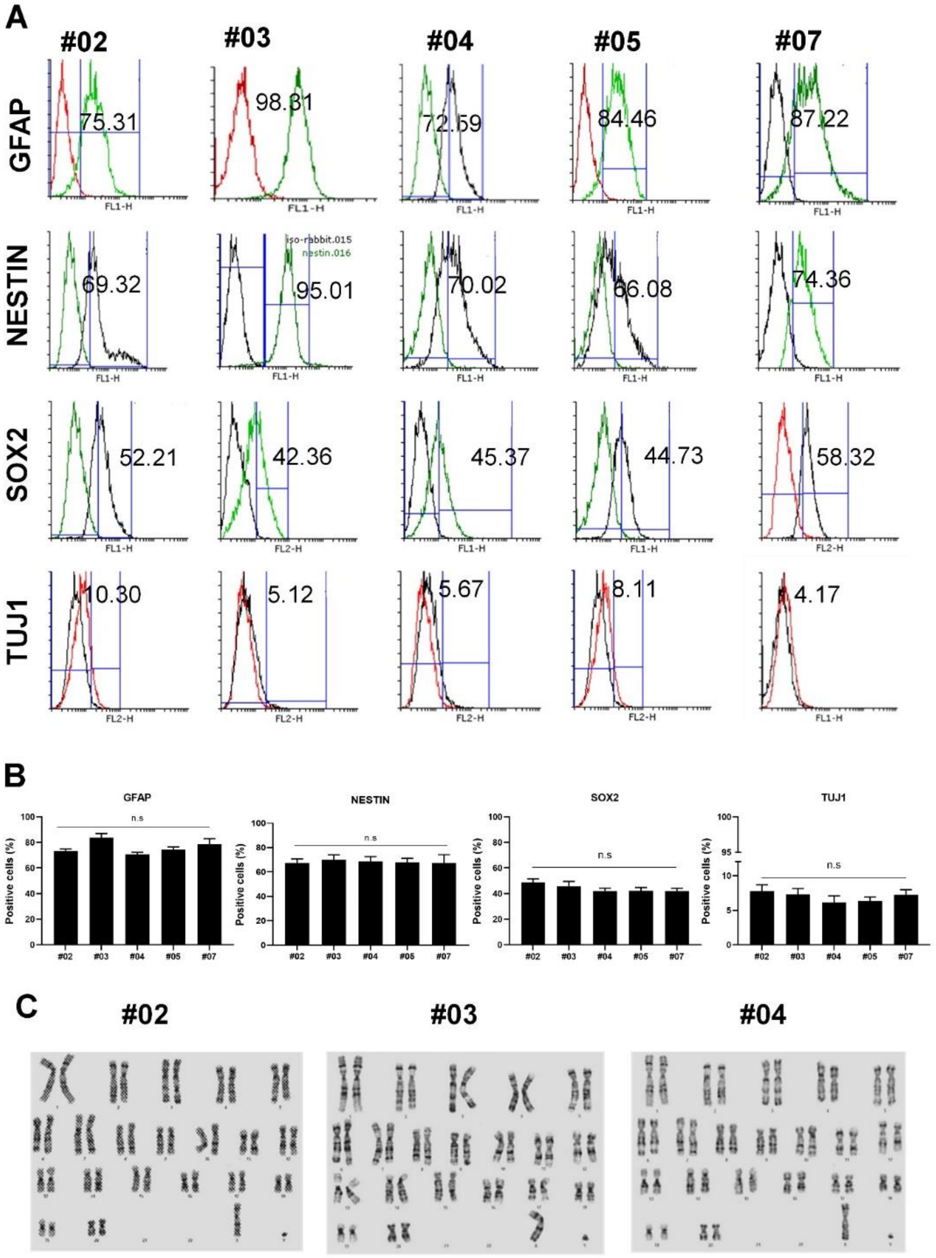


**Figure S2: fellow cytometry, Real-Time PCR analysis for individual monkeys, and Karyotype assessment**

**Fig. S-3**

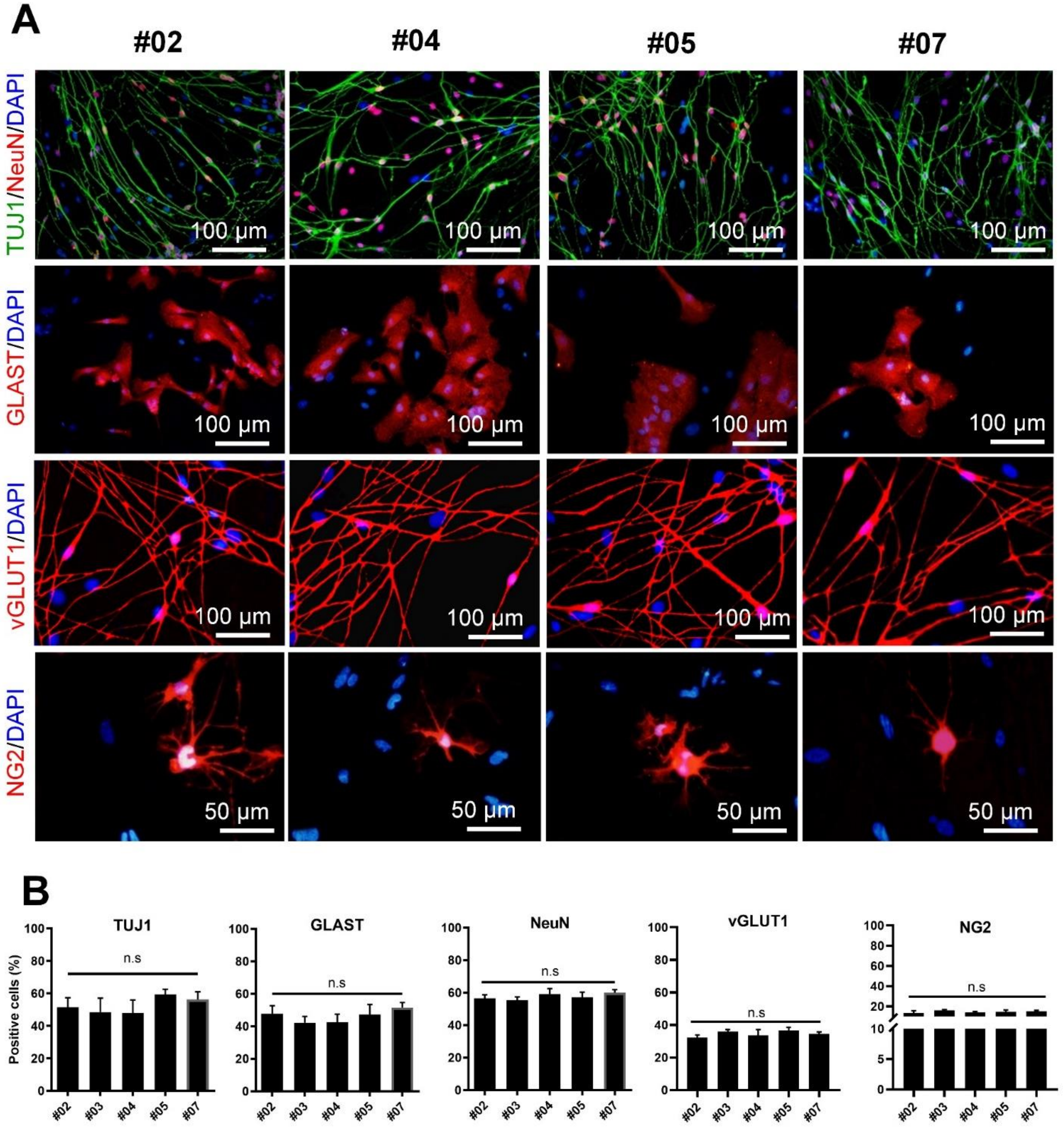


**Figure S3: spontaneous differentiation for isolated cells (Neuronal and glial markers), and quantification for different markers.**

#### Fig. S-4

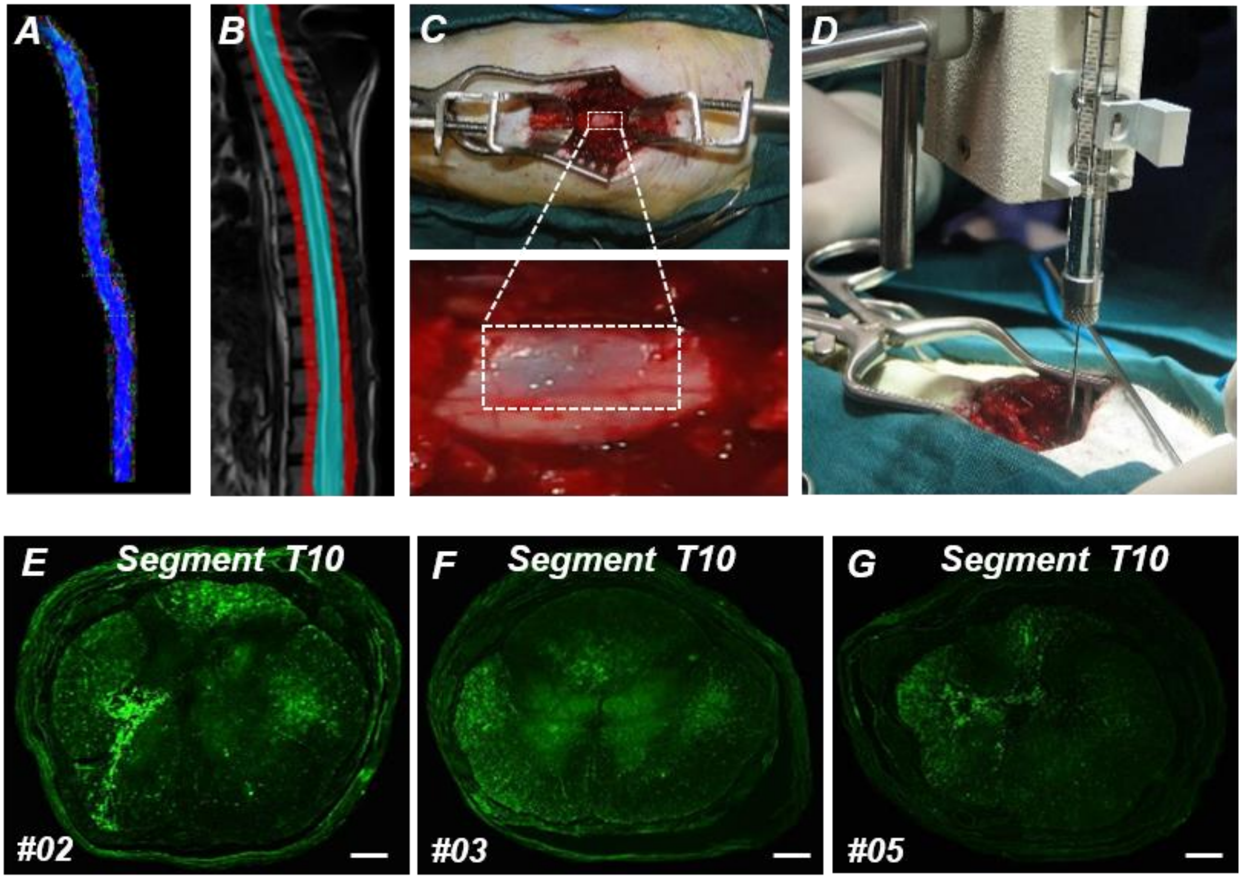


**Figure S4: tractography (MRI based), Spinal cord injury modeling and cell injection, and tracing the transplanted cells in monkeys**

**Fig. S-5**

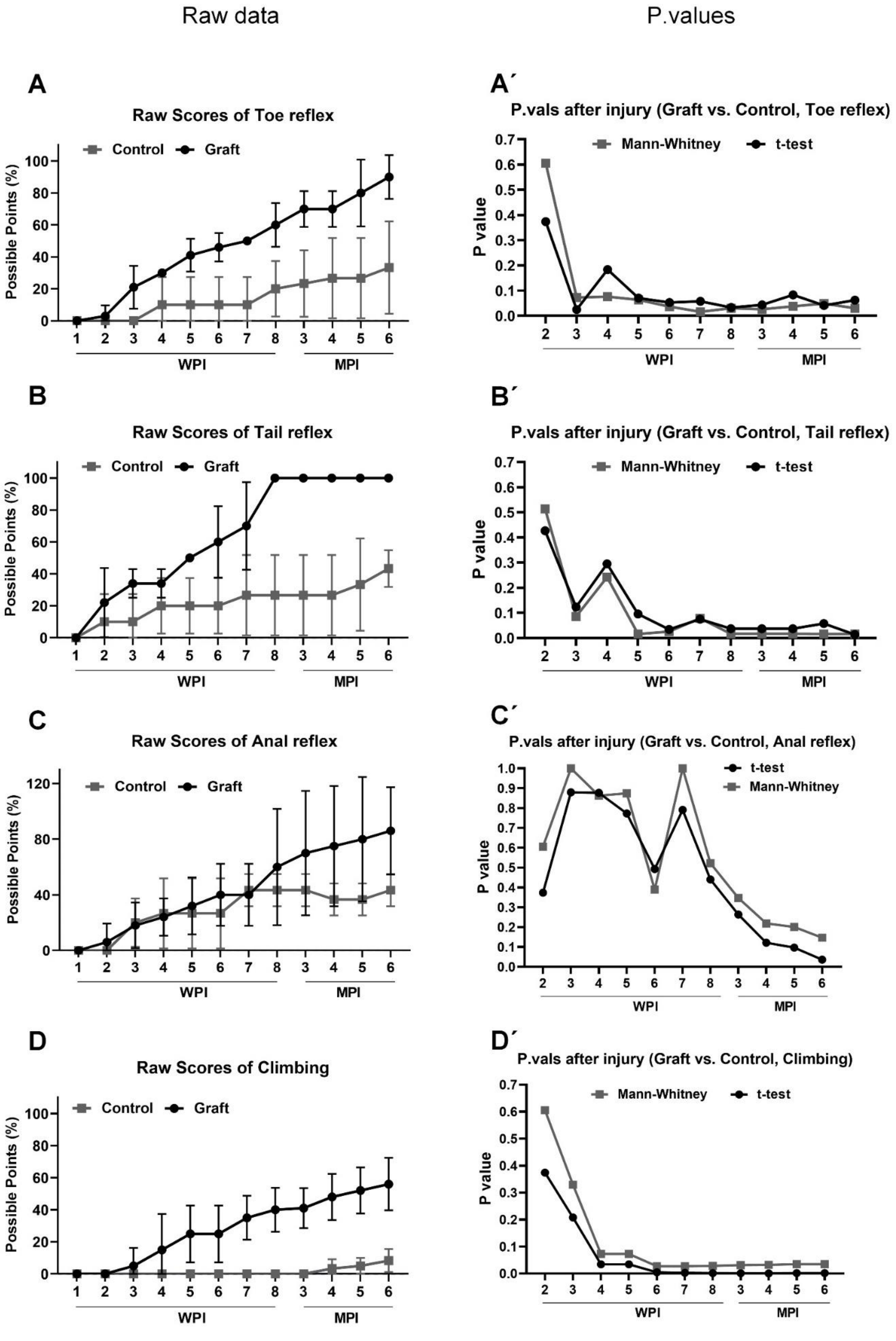


**Figure S5: Raw data and p value for sensory and motor function assessments**

#### Fig. S-6

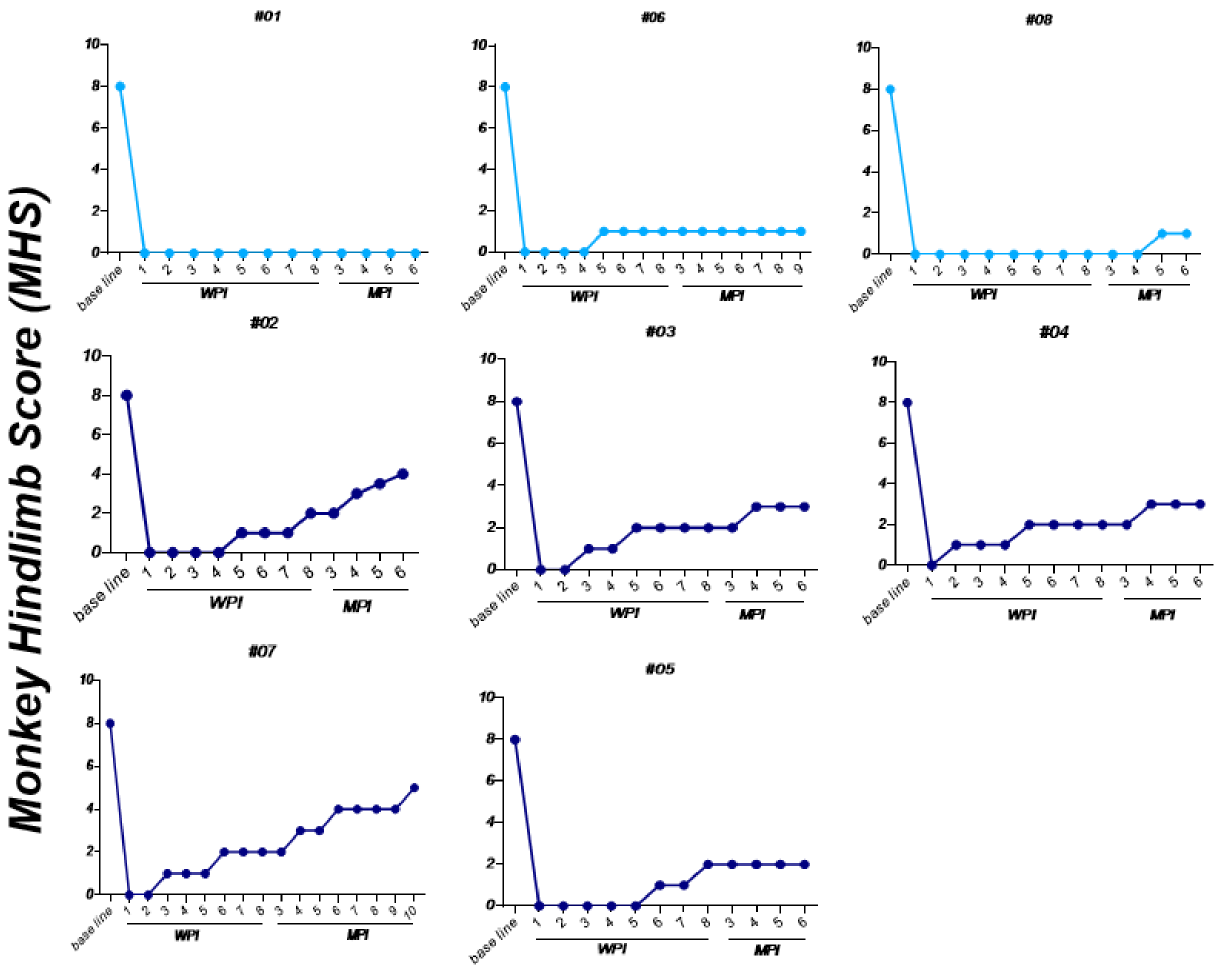


**Figure S6: Monkey hindlimb score for each animal**

Fig. S-7

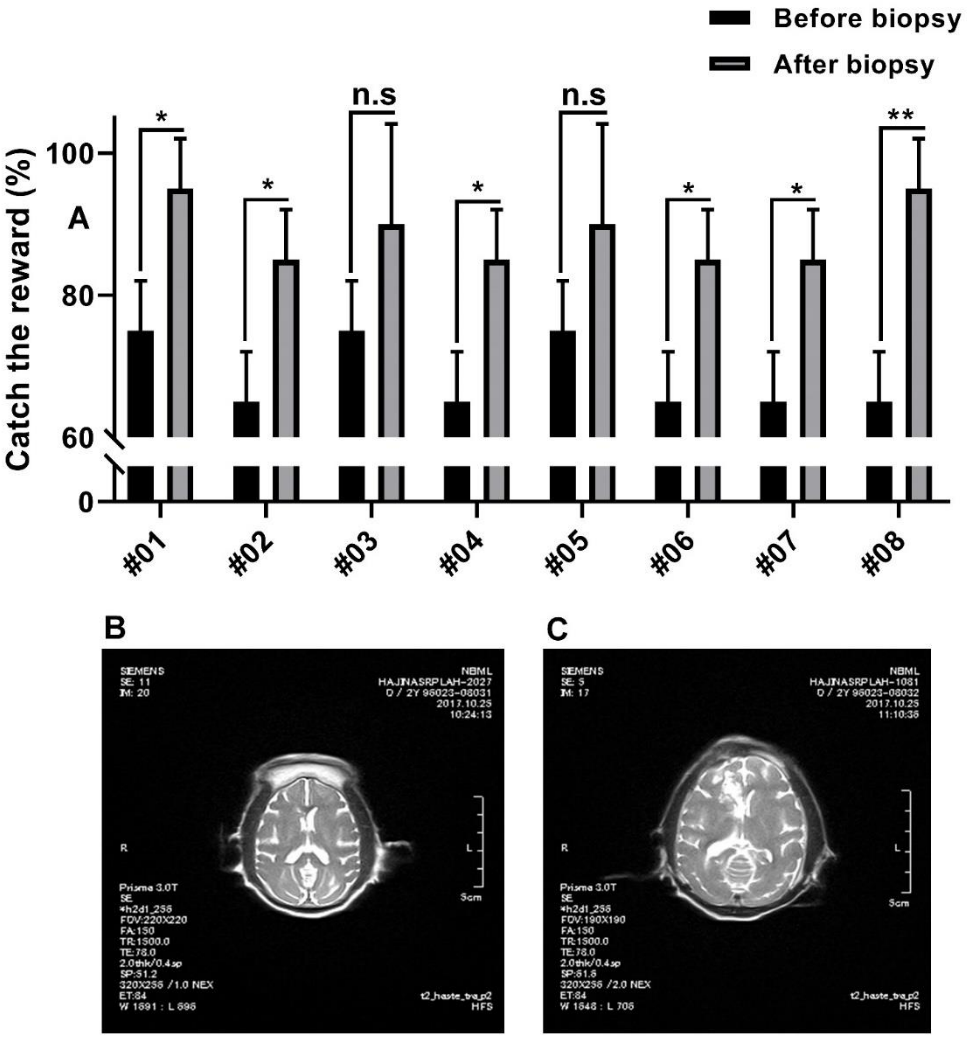


**Figure S7: response to reward before and after biopsy, and MR imaging after brain biopsy**

### Fig. S-8

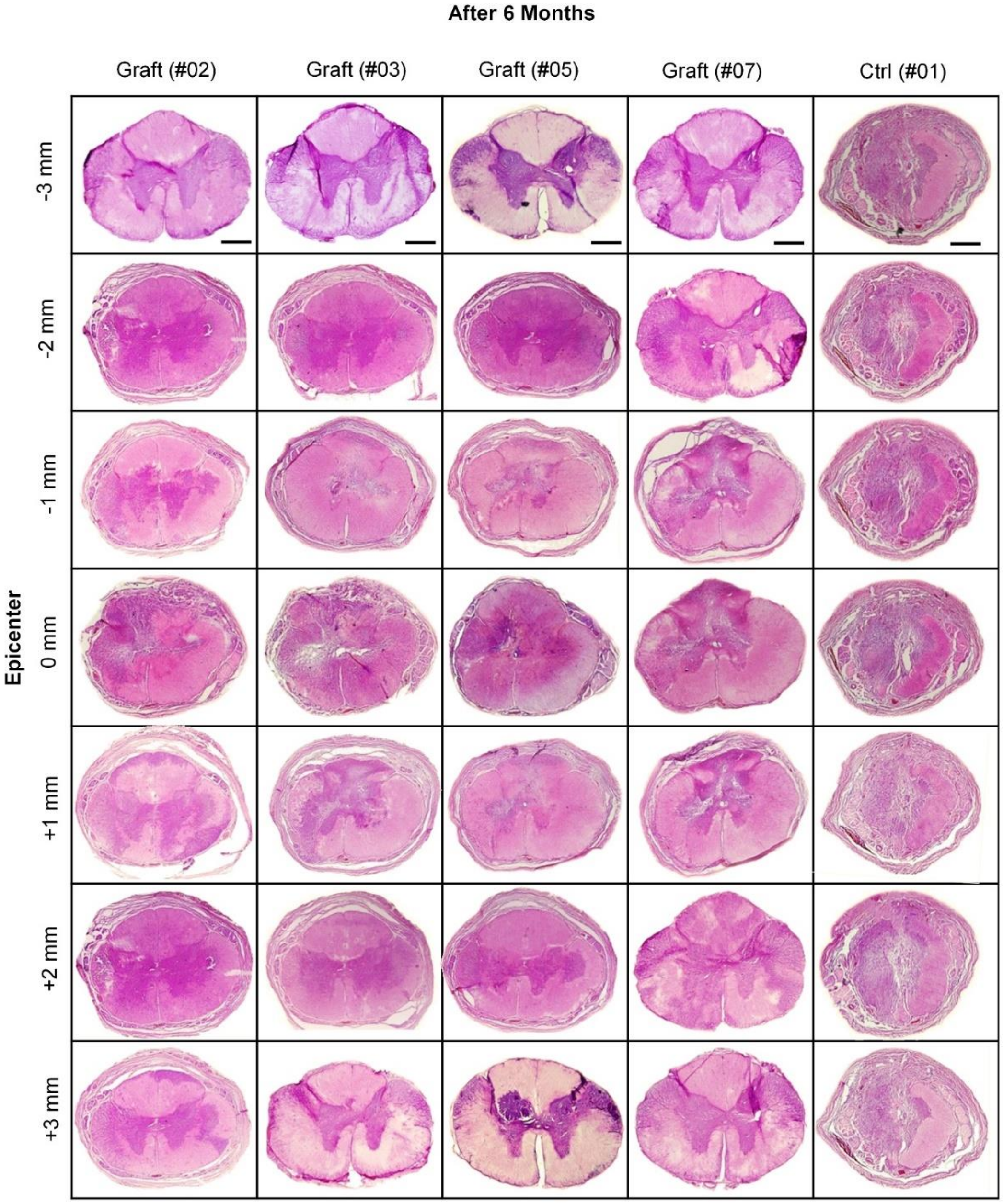


**Figure S8: H&E coronal sectioning**

### Fig. S-9

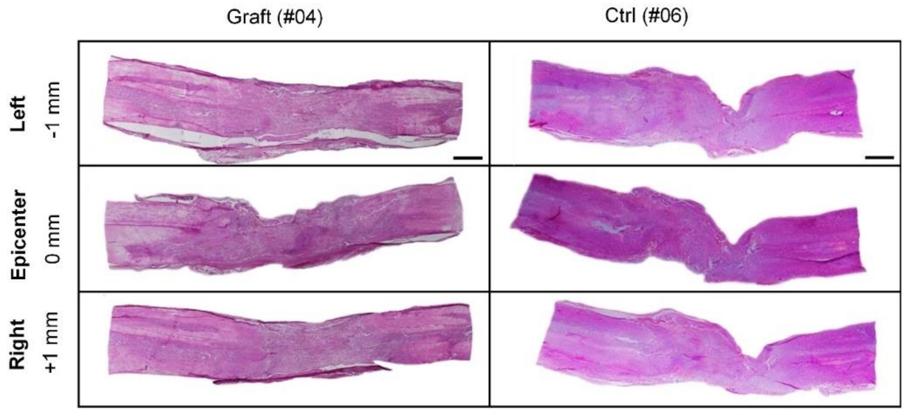


**Figure S9: H&E horizontal sectioning**

#### Fig. S-10

**ROYAN Institute**

Reproductive Biomedicine & Stem Cell Research Center

Number :EC/92/1009
May 05 ,2013

Dear Dr. Sahar Kiani

Your research project entitled "Autologous Neural Stem Cells Isolated from Sub-Ventricular Zone Can Improve Motor Abilities in Contusion Spinal Cord Injured Primate" has been discussed in Ethical Committee on May 05 ,2013 and has been approved under the following conditions:

1. Complete follow of National Codes of Research
2. Complete follow of Helsinki Declaration

This approval letter is issued just for the mentioned research title and has no other use.

Best regards

Shahverdi

Abdolhossein Shahverdi Ph.D. / Head of Ethical Committee

**Figure S10: Ethical approval letter**

| Table 2. Monkey hindlimb score (MHS) | | |
|---|---|---|
| | Score | |
| Corridor walking assay (walking ability as measured by hip, knee, and ankle movements) | Left | [illegible] |
| No voluntary movement in large joints (hip, knee or ankle) | 0 | 0 |
| Perceptible movement of 1–2 joints in the hindlimb | 1 | 1 |
| Perceptible movement of 3 joints | 2 | 2 |
| Vigorous movement of 3 joints without weight bearing | 3 | 3 |
| Occasional standing without sustainable weight bearing | 4 | 4 |
| Consistent weight support and able to walk with significant deficits | 5 | 5 |
| Standing and walking (3 m in N3 s) | 6 | 6 |
| Standing and walking (3 m in b3 s) | 7 | 7 |
| Coordinated walking forwards and backwards with intact body turning, jumping, and hopping | 8 | 8 |

| Table 3. Blood assessments | | Subject: #01 | |
|---|---|---|---|
| **Hematology** | | | |
| **CBC** | | | |
| **Test** | **Result** | **Unit** | **Normal range** |
| W.B.C | 11.8 | $\times10^3$/ul | 6-17.5 |
| R.B.C | 6.60 | Million/ul | 3.8-5.3 |
| Hemoglobin | 14.5 | g/dl | 10.5-14.5 |
| Hematocrit | 48.0 | % | 33.0-39.0 |
| M.C.V | 72.7 | fl | 70.0-86.0 |
| M.C.H | 22 | Pg | 23.0-31.0 |
| M.C.H.C | 30.2 | % | 30.0-36.0 |
| Platelets | 145.4 | $\times10^3$/ul | 145-450 |
| R.D.W | 14.7 | % | 11.0-15.0 |
| **Differential** | | | |
| Neutrophils | 40% | | |
| Lymphocyte | 50% | | |
| Monocyte | 9% | | |
| Eosinophil | 1% | | |
| **Biochemistry** | | | |
| **Test** | **Result** | **Unit** | **Normal range** |
| Fasting Serum Glucose | 45 | | |
| Urea | 40 | | |
| Creatinine | 0.8 | | |
| Sodium | 146 | Meq/L | 134-148 |
| Potassium | 5.21 | Meq/L | 3.5-5.5 |
| AST (SGOT) | 25 | U/L | Up to 32 |
| ALT (SGPT) | 15 | U/L | Up to 33 |
| **Immunology (ECLIA)** | | | |
| **Test** | **Result** | **Unit** | **Normal range** |
| Anti Cytomegalo Virus-IgG | 4.1 | IU/ml | Non-Reactive: <9.0 |
| Anti Cytomegalo Virus-IgM | Non-Reactive | Qual | Non-Reactive: <0.7 |
| Anti-HIV 1&2 Antibody | Non-Reactive | Qual | Non-Reactive |
| HBS Antigen | Non-Reactive | Qual | Non-Reactive |
| Anti-HBS Antibody | 0.1 | IU/L | Non-Reactive: <10 |
| Anti-HSV-1-IgG | Non-Reactive | Index | Negative: <0.9 |
| Anti-HSV-2-IgG | Non-Reactive | | Non-Reactive |
| Anti-HSV-1&2-IgM | Non-Reactive | IU/ml | Negative: <20 |
| **Subject: #02** | | | |
| **Hematology** | | | |
| **CBC** | | | |
| **Test** | **Result** | **Unit** | **Normal range** |
| W.B.C | 10.9 | $\times10^3$/ul | 6-17.5 |
| R.B.C | 5.80 | Million/ul | 3.8-5.3 |
| Hemoglobin | 13.3 | g/dl | 10.5-14.5 |
| Hematocrit | 43.5 | % | 33.0-39.0 |
| M.C.V | 75 | fl | 70.0-86.0 |
| M.C.H | 22.9 | Pg | 23.0-31.0 |
| M.C.H.C | 30.6 | % | 30.0-36.0 |
| Platelets | 375 | $\times10^3$/ul | 145-450 |
| R.D.W | 14.8 | % | 11.0-15.0 |
| **Differential** | | | |
| Neutrophils | 42% | | |
| Lymphocyte | 46% | | |
| Monocyte | 10% | | |
| Eosinophil | 2% | | |
| **Biochemistry** | | | |
| **Test** | **Result** | **Unit** | **Normal range** |
| Fasting Serum Glucose | 67 | | |
| Urea | 33 | | |
| Creatinine | 0.8 | | |
| Sodium | 145 | Meq/L | 134-148 |
| Potassium | 5.6 | Meq/L | 3.5-5.5 |

| | | | |
|---|---|---|---|
| AST (SGOT) | 22 | U/L | Up to 32 |
| ALT (SGPT) | 20 | U/L | Up to 33 |
| **Immunology (ECLIA)** | | | |
| **Test** | **Result** | **Unit** | **Normal range** |
| Anti Cytomegalo Virus-IgG | 1.3 | IU/ml | Non-Reactive: <9.0 |
| Anti Cytomegalo Virus-IgM | Non-Reactive | Qual | Non-Reactive: <0.7 |
| Anti-HIV 1&2 Antibody | Non-Reactive | Qual | Non-Reactive |
| HBS Antigen | Non-Reactive | Qual | Non-Reactive |
| Anti-HBS Antibody | 0.1 | IU/L | Non-Reactive: <10 |
| Anti-HSV-1-IgG | Non-Reactive | Index | Negative: <0.9 |
| Anti-HSV-2-IgG | Non-Reactive | | Non-Reactive |
| Anti-HSV-1&2-IgM | Non-Reactive | IU/ml | Negative: <20 |
| **Subject: #03** | | | |
| **Hematology** | | | |
| **CBC** | | | |
| **Test** | **Result** | **Unit** | **Normal range** |
| W.B.C | 10.5 | $\times 10^3$/ul | 6-17.5 |
| R.B.C | 5.65 | Million/ul | 3.8-5.3 |
| Hemoglobin | 14.2 | g/dl | 10.5-14.5 |
| Hematocrit | 47.5 | % | 33.0-39.0 |
| M.C.V | 75.8 | fl | 70.0-86.0 |
| M.C.H | 24 | Pg | 23.0-31.0 |
| M.C.H.C | 33.2 | % | 30.0-36.0 |
| Platelets | 235.3 | $\times 10^3$/ul | 145-450 |
| R.D.W | 12.4 | % | 11.0-15.0 |
| **Differential** | | | |
| Neutrophils | 48% | | |
| Lymphocyte | 46% | | |
| Monocyte | 5% | | |
| Eosinophil | 1% | | |
| **Biochemistry** | | | |
| **Test** | **Result** | **Unit** | **Normal range** |
| Fasting Serum Glucose | 58 | | |
| Urea | 37 | | |
| Creatinine | 0.7 | | |
| Sodium | 137 | Meq/L | 134-148 |
| Potassium | 4.30 | Meq/L | 3.5-5.5 |
| AST (SGOT) | 23 | U/L | Up to 32 |
| ALT (SGPT) | 18 | U/L | Up to 33 |
| **Immunology (ECLIA)** | | | |
| **Test** | **Result** | **Unit** | **Normal range** |
| Anti Cytomegalo Virus-IgG | 3.2 | IU/ml | Non-Reactive: <9.0 |
| Anti Cytomegalo Virus-IgM | Non-Reactive | Qual | Non-Reactive: <0.7 |
| Anti-HIV 1&2 Antibody | Non-Reactive | Qual | Non-Reactive |
| HBS Antigen | Non-Reactive | Qual | Non-Reactive |
| Anti-HBS Antibody | 0.1 | IU/L | Non-Reactive: <10 |
| Anti-HSV-1-IgG | Non-Reactive | Index | Negative: <0.9 |
| Anti-HSV-2-IgG | Non-Reactive | | Non-Reactive |
| Anti-HSV-1&2-IgM | Non-Reactive | IU/ml | Negative: <20 |
| **Subject: #04** | | | |
| **Hematology** | | | |
| **CBC** | | | |
| **Test** | **Result** | **Unit** | **Normal range** |
| W.B.C | 12.6 | $\times 10^3$/ul | 6-17.5 |
| R.B.C | 5.2 | Million/ul | 3.8-5.3 |
| Hemoglobin | 13.5 | g/dl | 10.5-14.5 |
| Hematocrit | 44.0 | % | 33.0-39.0 |
| M.C.V | 70.3 | fl | 70.0-86.0 |
| M.C.H | 25 | Pg | 23.0-31.0 |
| M.C.H.C | 33.25 | % | 30.0-36.0 |
| Platelets | 198.4 | $\times 10^3$/ul | 145-450 |
| R.D.W | 13.6 | % | 11.0-15.0 |
| **Differential** | | | |
| Neutrophils | 51% | | |

| | | | |
|---|---|---|---|
| Lymphocyte | 41% | | |
| Monocyte | 7% | | |
| Eosinophil | 1% | | |
| **Biochemistry** | | | |
| **Test** | **Result** | **Unit** | **Normal range** |
| Fasting Serum Glucose | 58 | | |
| Urea | 43 | | |
| Creatinine | 0.6 | | |
| Sodium | 132 | Meq/L | 134-148 |
| Potassium | 4.36 | Meq/L | 3.5-5.5 |
| AST (SGOT) | 20 | U/L | Up to 32 |
| ALT (SGPT) | 19 | U/L | Up to 33 |
| **Immunology (ECLIA)** | | | |
| **Test** | **Result** | **Unit** | **Normal range** |
| Anti Cytomegalo Virus-IgG | 3.0 | IU/ml | Non-Reactive: <9.0 |
| Anti Cytomegalo Virus-IgM | Non-Reactive | Qual | Non-Reactive: <0.7 |
| Anti-HIV 1&2 Antibody | Non-Reactive | Qual | Non-Reactive |
| HBS Antigen | Non-Reactive | Qual | Non-Reactive |
| Anti-HBS Antibody | 0.1 | IU/L | Non-Reactive: <10 |
| Anti-HSV-1-IgG | Non-Reactive | Index | Negative: <0.9 |
| Anti-HSV-2-IgG | Non-Reactive | | Non-Reactive |
| Anti-HSV-1&2-IgM | Non-Reactive | IU/ml | Negative: <20 |
| **Subject: #05** | | | |
| **Hematology** | | | |
| **CBC** | | | |
| **Test** | **Result** | **Unit** | **Normal range** |
| W.B.C | 13.8 | $\times 10^3$/ul | 6-17.5 |
| R.B.C | 4.1 | Million/ul | 3.8-5.3 |
| Hemoglobin | 13.7 | g/dl | 10.5-14.5 |
| Hematocrit | 38.5 | % | 33.0-39.0 |
| M.C.V | 72.3 | fl | 70.0-86.0 |
| M.C.H | 24 | Pg | 23.0-31.0 |
| M.C.H.C | 34.2 | % | 30.0-36.0 |
| Platelets | 211.3 | $\times 10^3$/ul | 145-450 |
| R.D.W | 12.6 | % | 11.0-15.0 |
| **Differential** | | | |
| Neutrophils | 48% | | |
| Lymphocyte | 43% | | |
| Monocyte | 7% | | |
| Eosinophil | 2% | | |
| **Biochemistry** | | | |
| **Test** | **Result** | **Unit** | **Normal range** |
| Fasting Serum Glucose | 73 | | |
| Urea | 38 | | |
| Creatinine | 1.1 | | |
| Sodium | 126 | Meq/L | 134-148 |
| Potassium | 3.9 | Meq/L | 3.5-5.5 |
| AST (SGOT) | 23 | U/L | Up to 32 |
| ALT (SGPT) | 21 | U/L | Up to 33 |
| **Immunology (ECLIA)** | | | |
| **Test** | **Result** | **Unit** | **Normal range** |
| Anti Cytomegalo Virus-IgG | 2.5 | IU/ml | Non-Reactive: <9.0 |
| Anti Cytomegalo Virus-IgM | Non-Reactive | Qual | Non-Reactive: <0.7 |
| Anti-HIV 1&2 Antibody | Non-Reactive | Qual | Non-Reactive |
| HBS Antigen | Non-Reactive | Qual | Non-Reactive |
| Anti-HBS Antibody | 1.1 | IU/L | Non-Reactive: <10 |
| Anti-HSV-1-IgG | Non-Reactive | Index | Negative: <0.9 |
| Anti-HSV-2-IgG | Non-Reactive | | Non-Reactive |
| Anti-HSV-1&2-IgM | Non-Reactive | IU/ml | Negative: <20 |
| **Subject: #06** | | | |
| **Hematology** | | | |
| **CBC** | | | |
| **Test** | **Result** | **Unit** | **Normal range** |
| W.B.C | 10.3 | $\times 10^3$/ul | 6-17.5 |

| | | | |
|---|---|---|---|
| R.B.C | 4.1 | Million/ul | 3.8-5.3 |
| Hemoglobin | 12.5 | g/dl | 10.5-14.5 |
| Hematocrit | 35.0 | % | 33.0-39.0 |
| M.C.V | 73.8 | fl | 70.0-86.0 |
| M.C.H | 25 | Pg | 23.0-31.0 |
| M.C.H.C | 32.0 | % | 30.0-36.0 |
| Platelets | 245.4 | ×$10^3$/ul | 145-450 |
| R.D.W | 12.7 | % | 11.0-15.0 |
| **Differential** | | | |
| Neutrophils | 50% | | |
| Lymphocyte | 42% | | |
| Monocyte | 6% | | |
| Eosinophil | 2% | | |
| **Biochemistry** | | | |
| **Test** | **Result** | **Unit** | **Normal range** |
| Fasting Serum Glucose | 68 | | |
| Urea | 40 | | |
| Creatinine | 0.9 | | |
| Sodium | 136 | Meq/L | 134-148 |
| Potassium | 4.2 | Meq/L | 3.5-5.5 |
| AST (SGOT) | 20 | U/L | Up to 32 |
| ALT (SGPT) | 23 | U/L | Up to 33 |
| **Immunology (ECLIA)** | | | |
| **Test** | **Result** | **Unit** | **Normal range** |
| Anti Cytomegalo Virus-IgG | 2.2 | IU/ml | Non-Reactive: <9.0 |
| Anti Cytomegalo Virus-IgM | Non-Reactive | Qual | Non-Reactive: <0.7 |
| Anti-HIV 1&2 Antibody | Non-Reactive | Qual | Non-Reactive |
| HBS Antigen | Non-Reactive | Qual | Non-Reactive |
| Anti-HBS Antibody | 0.7 | IU/L | Non-Reactive: <10 |
| Anti-HSV-1-IgG | Non-Reactive | Index | Negative: <0.9 |
| Anti-HSV-2-IgG | Non-Reactive | | Non-Reactive |
| Anti-HSV-1&2-IgM | Non-Reactive | IU/ml | Negative: <20 |
| **Subject: #07** | | | |
| **Hematology** | | | |
| **CBC** | | | |
| **Test** | **Result** | **Unit** | **Normal range** |
| W.B.C | 11.7 | ×$10^3$/ul | 6-17.5 |
| R.B.C | 4.4 | Million/ul | 3.8-5.3 |
| Hemoglobin | 11.5 | g/dl | 10.5-14.5 |
| Hematocrit | 37.2 | % | 33.0-39.0 |
| M.C.V | 76.8 | fl | 70.0-86.0 |
| M.C.H | 24 | Pg | 23.0-31.0 |
| M.C.H.C | 32.3 | % | 30.0-36.0 |
| Platelets | 185.1 | ×$10^3$/ul | 145-450 |
| R.D.W | 12.8 | % | 11.0-15.0 |
| **Differential** | | | |
| Neutrophils | 54% | | |
| Lymphocyte | 40% | | |
| Monocyte | 5% | | |
| Eosinophil | 1% | | |
| **Biochemistry** | | | |
| **Test** | **Result** | **Unit** | **Normal range** |
| Fasting Serum Glucose | 63 | | |
| Urea | 42 | | |
| Creatinine | 0.5 | | |
| Sodium | 128 | Meq/L | 134-148 |
| Potassium | 4.23 | Meq/L | 3.5-5.5 |
| AST (SGOT) | 23 | U/L | Up to 32 |
| ALT (SGPT) | 17 | U/L | Up to 33 |
| **Immunology (ECLIA)** | | | |
| **Test** | **Result** | **Unit** | **Normal range** |
| Anti Cytomegalo Virus-IgG | 4.2 | IU/ml | Non-Reactive: <9.0 |
| Anti Cytomegalo Virus-IgM | Non-Reactive | Qual | Non-Reactive: <0.7 |
| Anti-HIV 1&2 Antibody | Non-Reactive | Qual | Non-Reactive |
| HBS Antigen | Non-Reactive | Qual | Non-Reactive |

| | | | |
|---|---|---|---|
| Anti-HBS Antibody | 0.1 | IU/L | Non-Reactive: <10 |
| Anti-HSV-1-IgG | Non-Reactive | Index | Negative: <0.9 |
| Anti-HSV-2-IgG | Non-Reactive | | Non-Reactive |
| Anti-HSV-1&2-IgM | Non-Reactive | IU/ml | Negative: <20 |
| **Subject: #08** | | | |
| **Hematology** | | | |
| **CBC** | | | |
| **Test** | **Result** | **Unit** | **Normal range** |
| W.B.C | 10.8 | $\times 10^3$/ul | 6-17.5 |
| R.B.C | 4.9 | Million/ul | 3.8-5.3 |
| Hemoglobin | 12.1 | g/dl | 10.5-14.5 |
| Hematocrit | 35.0 | % | 33.0-39.0 |
| M.C.V | 76.5 | fl | 70.0-86.0 |
| M.C.H | 28 | Pg | 23.0-31.0 |
| M.C.H.C | 31.23 | % | 30.0-36.0 |
| Platelets | 215.7 | $\times 10^3$/ul | 145-450 |
| R.D.W | 12.2 | % | 11.0-15.0 |
| **Differential** | | | |
| Neutrophils | 53% | | |
| Lymphocyte | 41% | | |
| Monocyte | 4% | | |
| Eosinophil | 2% | | |
| **Biochemistry** | | | |
| **Test** | **Result** | **Unit** | **Normal range** |
| Fasting Serum Glucose | 76 | | |
| Urea | 41 | | |
| Creatinine | 0.8 | | |
| Sodium | 124 | Meq/L | 134-148 |
| Potassium | 4.36 | Meq/L | 3.5-5.5 |
| AST (SGOT) | 21 | U/L | Up to 32 |
| ALT (SGPT) | 17 | U/L | Up to 33 |
| **Immunology (ECLIA)** | | | |
| **Test** | **Result** | **Unit** | **Normal range** |
| Anti Cytomegalo Virus-IgG | 4.0 | IU/ml | Non-Reactive: <9.0 |
| Anti Cytomegalo Virus-IgM | Non-Reactive | Qual | Non-Reactive: <0.7 |
| Anti-HIV 1&2 Antibody | Non-Reactive | Qual | Non-Reactive |
| HBS Antigen | Non-Reactive | Qual | Non-Reactive |
| Anti-HBS Antibody | 0.2 | IU/L | Non-Reactive: <10 |
| Anti-HSV-1-IgG | Non-Reactive | Index | Negative: <0.9 |
| Anti-HSV-2-IgG | Non-Reactive | | Non-Reactive |
| Anti-HSV-1&2-IgM | Non-Reactive | IU/ml | Negative: <20 |
